# High rate of mutation and efficient removal by selection of structural variants from natural populations of *Caenorhabditis elegans*

**DOI:** 10.1101/2025.03.22.644739

**Authors:** Ayush Shekhar Saxena, Charles F. Baer

## Abstract

The importance of genomic structural variants (SVs) is well-appreciated, but less is known about their mutational properties than of single nucleotide variants (SNVs) and short indels. The reason is simple: the longer the variant, the less likely it will be covered by a single sequencing read, thus the harder it is to map unambiguously to a unique genomic location.

Here we report SV mutation rate estimates from six mutation accumulation (MA) lines from two strains of *C. elegans* using long-read (PacBio) sequencing. The inferred SV mutation rate is ∼0.03/genome/generation, about 1/10 the SNV rate and 1/4 the short indel rate. We identified 40 SV mutations (12 insertions, 28 deletions, 0 inversions) and 52 false positive (FP) variants by manual inspection. Excluding one atypical line (5 mutations, 35 FPs), the signal (mutant) to noise (FP) ratio is approximately 2:1. False negative rates were determined by simulating variants in the reference genome and observing ’recall’. Recall rate ranges from >90% for short indels and declines as SV length increases. Small deletions have nearly the same recall rate as small insertions (∼100bp), but deletions have higher recall rates than insertions as size increases. The reported SV mutation rate is likely a lower bound.

A quarter of identified SV mutations occur in SV hotspots that harbor pre-existing low complexity repeat variation. Comparison of the spectrum of spontaneous SVs to wild isolates implies that natural selection is not only efficient at removing SVs in exons but also effectively removes SVs in intergenic regions.

## Introduction

Based on the total number of bases affected, structural variants (SV) — large insertions and deletions, repeat expansions and contractions, inversions, and translocations — are the largest source of genetic diversity in the genomes of multicellular organisms, and they have the largest average phenotypic and fitness effects of any type of mutation (Collins et al. 2020; Beyter et al. 2021). SVs are associated with a host of human disorders, including autism (Marshall et al. 2008; Weiss et al. 2008; Pinto et al. 2010; Sanders et al. 2011), schizophrenia (Walsh et al. 2008; Karayiorgou et al. 2010), and several types of cancers (Friedman et al. 1994; Wooster et al. 1995; Meyerson and Pellman 2011; Stransky et al. 2014; Nattestad et al. 2018). The functional consequences of SVs on phenotype and fitness are well-documented (Hurles et al. 2008; Conrad et al. 2010; Yi and Ju 2018; Loegler et al. 2025).

In recent years, human population geneticists have converged on the idea that the genetic basis of complex disease can be largely accounted for by the combined influence of very rare, recently arisen variants of large effect (Wang et al. 2021) and variants of small effect at very many loci (the “omnigenome”, Boyle et al. 2017). Both classes of variants imply a significant role for mutation, the former for obvious reasons and the latter because, even if the per-locus mutation rate is low, the cumulative genetic variance contributed by a large fraction of the genome at mutation-selection-drift balance may be substantial.

Given the outsized contribution of SVs to genetic diversity on the one hand, and the apparently important contribution of mutation to complex disease on the other, understanding the mutational properties of SVs is an important problem in biomedical genetics. More broadly, characterizing SV mutation is necessary for an unbiased estimate of the distribution of fitness effects (DFE) of new mutations, which is a longstanding goal of evolutionary genetics (Fisher 1922; Eyre-Walker and Keightley 2007). Nearly all estimates of the DFE are restricted to the effects of SNPs and small indels (and most are restricted to SNPs, e.g., Kousathanas and Keightley 2013; Kim et al. 2017; Tataru et al. 2017; Böndel et al. 2019; Gilbert et al. 2021; James et al. 2023; Crombie et al. 2024). Because SVs are ignored, all the variation in fitness is ascribed to small indels and SNPs, artificially inflating their importance by an unknown amount that may be quite large.

The reason that the mutational properties of SVs are poorly characterized is not because their biological relevance is not appreciated; it obviously is. Rather, the problem is technical: SVs are challenging to unambiguously identify and characterize with short-read sequencing technology, which remains the dominant mode of whole-genome sequencing. Long-read sequencing is superior to short-read sequencing for SV calling for several reasons. Longer reads may span the entire region of interest in one contiguous molecule, including problematic regions such as low complexity repeats. Even when a single read does not span the entire region of interest, longer reads result in superior local *de novo* assemblies that can span regions much larger than the read length itself. The obvious disadvantages include cost and base-calling error rate with respect to base-substitution and small indel calls, although the accuracy of the latest generation of some long-read platforms approaches that of short-read sequencing (Harvey et al. 2023).

A complete *de novo* assembly (whole genome assembly, WGA) of an individual is the only method certain to capture the full array of structural variants in a genome (Chaisson et al. 2015). As the cost of long-read technology declines, genome assemblies from single individuals have become economically feasible. Two recent studies highlight the rapidity and magnitude of advances in the field. Loegler et al. (2025) reported the genotype to phenotype relationship of >1000 yeast strains based on long-read sequencing. They found that including SVs (and small indels) in the analyses increased the average explained heritability of >8000 traits by ∼14% above that explained by SNPs alone. Schloissnig et al. (2025) estimated the frequency of SVs in >1000 human genomes, resulting in a ∼4X increase in the estimated frequency of SVs in the average human genome. However, the application of long-read sequencing technology to characterize spontaneous mutations is still nascent, although it is evident that long-read sequencing picks up de novo variants that are missed with short-read sequencing (Noyes et al. 2022; Lopez-Cortegano et al. 2023; Zhou et al. 2023; Hénault et al. 2024; López-Cortegano et al. 2025; Porubsky et al. 2025).

Here we present estimates of the spontaneous mutation rate and spectrum of SVs in a set of *C. elegans* mutation accumulation (MA) lines propagated under minimal selection for ∼250 generations. A schematic diagram of the MA experiment is depicted in Figure 1. Four MA lines derived from the N2 strain and two MA lines derived from the PB306 strain, and their ancestral progenitors, were sequenced using Pacific Biosciences Sequel and Sequel 2 technology. In addition, we report estimates of the standing SV diversity derived from long-read sequences of the genomes of four wild isolates of *C. elegans*. Comparison of the relative proportions of SVs to other variants (SNPs and small indels) among spontaneous mutations to the same proportions in the standing variation provides an estimate of the relative strength of selection against SVs in nature.

**Figure 1.**
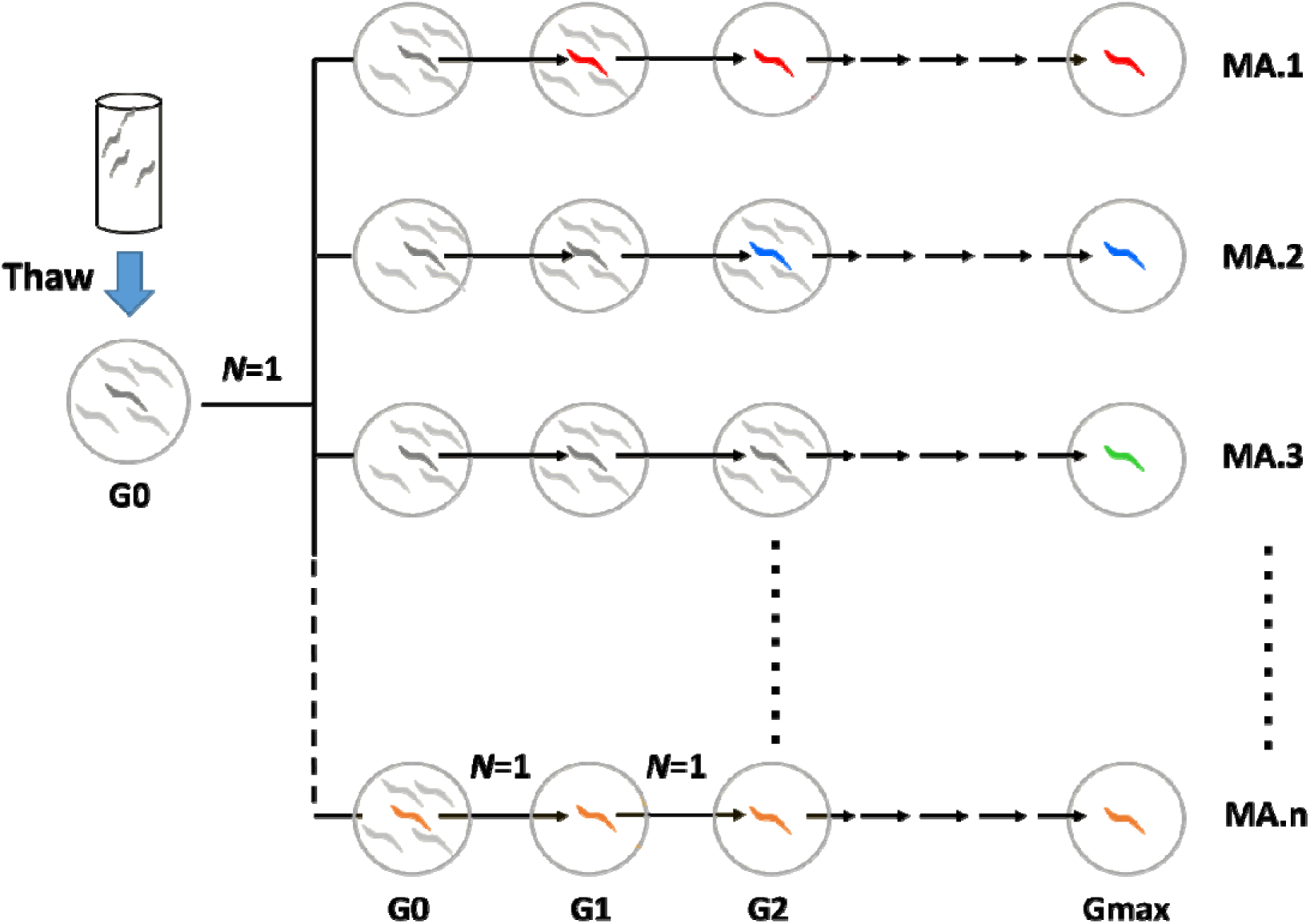
Schematic diagram of the mutation accumulation (MA) experiment. A cryopreserved sample of the ancestor (N2 or PB306) was thawed onto an agar plate (G0, left) and grown for a few days until the cryopreserved worms had reached the L4 larval stage. L4-stage worms were transferred singly to new NGM agar plates; those were the founders of the MA lines (MA.1, MA.2,…MA.n; n=100). At four-day intervals (one generation), a single immature worm was picked at random onto a new plate. This procedure was repeated until the experiment reached Gmax=250 generations. Individual MA lines experienced fewer than Gmax generations, Gave=238 generations. The different colored worms represent (the set of) unique mutations fixed in each MA line, which contribute to the among-line (mutational) variance.

## Results

### Workflow

We define a structural variant (SV) as a variant larger than 30 base pairs (bp), using a combination of alignment and assembly, followed by conventional variant calling. We chose 30 bp as our lower bound for designating a variant as an SV rather than the more common 50 bp because preliminary analysis revealed that long-read sequencing allowed us to detect indels between 30-50 bp that were missed by short-read sequencing. The difference is only meaningful when it comes to comparison of SV rates between this study and studies that use 50 bp as the lower bound; we note the discrepancy where appropriate.

Recognizing the inherent challenges in SV detection, we attempted to automate the process of discovering the optimal parameters for the variant calling pipeline using segregating SVs as ‘ground truth’, under the assumption that a variant called across many wild isolates is likely real. We discovered that the signal-to-noise ratio of spontaneous SVs is less than that of segregating SVs, complicating the development of an appropriate pipeline. These complications are detailed in section 2 of the Supplementary Discussion. We ultimately adopted a workflow loosely based on the methodology of Audano et al. (2019), with three key modifications:

1. Utilization of a superior aligner, minimap2 (Li 2018), instead of blasr (https://github.com/jcombs1/blasr);
2. Variant calling with PBSV (https://github.com/PacificBiosciences/pbsv); and, critically,
3. Manual (“by eye”) verification of each SV call by inspection in IGV (Robinson et al. 2017).

This workflow involves aligning error-prone PacBio raw reads (‘subreads’) to the *C. elegans* reference genome, followed by *de novo* assembly of aligned reads in 60-kb tiles. Base calling errors in raw subreads can exceed 10% (Weirather et al. 2017; Zhang et al. 2020). The assembled contigs — henceforth referred to as “pseudo-reads”— are then re-aligned to the reference genome, followed by conventional variant calling using PBSV. The pseudo-reads exhibit higher sequence quality than raw PacBio subreads, and aligned pseudo-reads are free from alignment errors that may occur with subread alignment. In a conventional SV-calling workflow, variants are genotyped only in the sample relative to the reference genome. In contrast, identifying *de novo* (spontaneous) SVs requires accurate genotype calls in <u>both</u> the MA progenitor and the derived MA line. Because false positives can arise from genotyping error in either sample (parent or MA), the spontaneous-mutation variant calling workflow inherently carries a higher false-positive rate than conventional SV calling. Consequently, each final mutation call is manually verified using IGV.

Supplementary Table 1 summarizes the base-calling error rates of the raw PacBio subreads and the assembly-derived pseudo-reads. The error rate of raw reads, calculated from all N2-derived lines, is 14%, while the pseudo-reads exhibit a significantly reduced per base error rate of 2.2%. Base-call error rates were calculated after alignment using the ’NM’ tag in the BAM file (See Methods), which represents the Levenshtein distance between the aligned segment of the read and the reference. Notably, pseudo-reads also exhibit fewer unaligned segments, measured as soft or hard clips after alignment. The percentage of clipped bases upon alignment is approximately 5.3% for raw reads, compared to 4.3% for pseudo-reads. To minimize the effect of reference bias on base-call error rate estimates, we only report results from N2-derived lines, from which the reference genome is derived, thereby providing more reliable estimates. The gains to alignment quality are even higher for wild isolates and PB306-derived MA lines. Base-calling error rates and clipping percentages for the remaining lines are likewise presented in Supplementary Table 1.

Raw sequence data generated in this study are archived in the NCBI Short Read Archive (PRJNA1233366). We also utilized publicly available long-read data of *C. elegans* wild isolates from the following SRA repositories (PRJNA523481, and PRJNA1025857).

### SV mutation rate

The mutation rate estimated from observed SVs (i.e., not accounting for false negatives) is 0.03/genome/generation (7 SVs per MA line, over 238 generations per MA line, on average). In comparison, we identified an average of 62 single nucleotide variants (SNV) and 25 short indel mutations per line in the same set of lines. The inferred SV mutation rate is roughly 10% of the SNV rate and 30% of the short indel rate, indicating that SVs comprise about 7.5% of new mutations (6% by the more traditional >50 bp criterion), or roughly one new SV mutation per-genome every ∼30 generations. Summary statistics of SV mutations are presented in Table 1. The full set of SVs and their attributes are given in Supplementary Tables 2 (N2) and 3 (PB306).

**Table 1.**
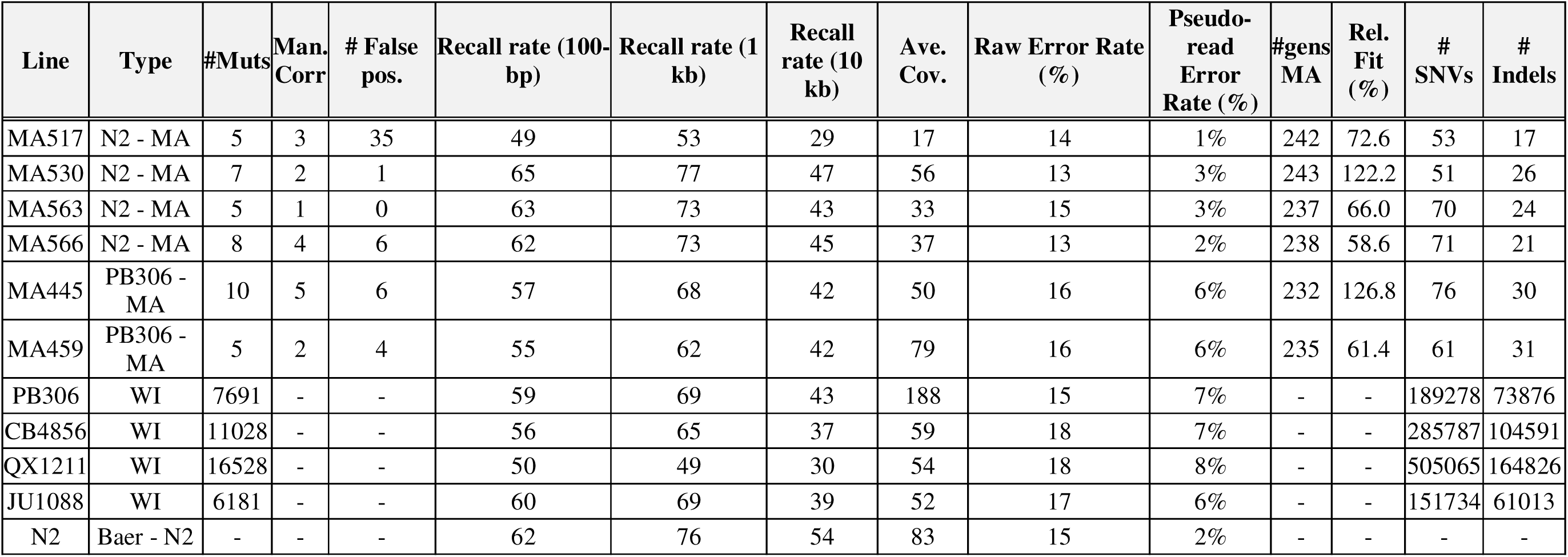
Summary statistics. . Column headings: *Line* (strain.MA line # or wild isolate ID); *Type* (MA or Wild isolate); *#Muts* (number of inferred structural variants > 30bp in size); *Man. Corr* (# of SVs that required manual correction; see Methods for explanation); #*False pos.* (# of false positives); *%Recall (100 bp, 1Kb, 10Kb)* (% of simulated SVs of that length recalled; see Methods for explanation); *Ave. Cov*. (average sequencing coverage); *Raw error rate* (Error rate of raw PacBio subreads calculated after alignment to reference); *Pseudo-reads Error rate* (Error rate of assembled contigs after alignment to reference)*; #gens MA* (number of MA generations; see Methods); *Rel. fit (%)* (Fitness of MA lines relative to ancestor N2; unpublished data from Saxena et al.); #SNVs (number of base-substitution mutations); #Indels (number of indel mutations ≤ 30 bp). “Mutations” in wild isolates are variant compared to the N2 reference genome. For a more detailed discussion of false positives in wild isolates, see section 2 of Supplementary Discussion.

### Estimation of False Discovery Rates

#### (a) False positives

The accuracy of SV mutation rate estimates is contingent upon the accuracy of the variant calling workflow. False positives can be identified only by either (*i*) “by eye” validation of putative variants using Integrative Genomics Viewer (IGV), or (*ii*) by using a different sequencing technique, ideally by PCR amplification of the putative mutant locus followed by Sanger sequencing of the resulting amplicon. However, those approaches become impractical as the number of samples and/or putative variants increases.

We scrutinized each putative variant in the MA lines by visualizing the aligned pseudo-reads, raw PacBio subreads, and Illumina reads from a separate study in IGV. We identified 52 false positives in the original set of called putative SVs, resulting in a signal (inferred mutant, n=40) to noise (false positive, n=52) ratio of 40/52 = 0.77. However, 2/3 of the false positives occurred in one MA line (N2 line MA517). MA517 had lower sequencing coverage than the other lines (17X vs an average of 51X for other MA lines), which potentially introduced alignment and/or assembly artifacts. When line MA517 is excluded, the signal-to-noise ratio increases to ∼2:1.

Among the true positive calls, minor corrections were made after visualizing the SVs in IGV. Corrections are required when an MA SV occurs at a locus that already carries an ancestral SV. Including other minor reasons for adjustments, corrections were necessary for 17 of the 40 true positive calls. The reasons for corrections are provided in Supplementary Tables 2 and 3.

**Table 2.** SV spectrum by variant type and number of bases affected.

| Line | Type | Insertions | Deletions | Inversions | Net bases affected | Net base gain (loss) |
| --- | --- | --- | --- | --- | --- | --- |
| N2.517 | MA | 2 | 3 | 0 | 5,342 | 184 |
| N2.530 | MA | 3 | 4 | 0 | 1,509 | 843 |
| N2.563 | MA | 2 | 3 | 0 | 7,606 | (309) |
| N2.566 | MA | 0 | 8 | 0 | 2,166 | (2,166) |
| PB.445 | MA | 3 | 7 | 0 | 3,501 | (1,397) |
| PB.459 | MA | 2 | 3 | 0 | 12,184 | 8,536 |
| PB306 | WI | 4347 | 3280 | 64 | 4,027,284 | 1,791,544 |
| CB4856 | WI | 5887 | 5056 | 85 | 5,205,431 | 1,562,847 |
| QX1211 | WI | 9148 | 7279 | 101 | 7,949,707 | 2,849,281 |
| JU1088 | WI | 3597 | 2525 | 59 | 3,331,096 | 1,465,676 |

**Table 3.** Distribution of variant types by genomic context. See Equation 1 in the Methods for details of calculation of relative diversity *w*\*. The subscripts “*All*” and “*PB306*” for *w*\* and *p*-values represent values calculated from all four wild isolates and PB306 only, respectively. Values of *w\** in bold font are statistically significant (׈ = 0.05); “*p”* is the *p*-value of the 2x2 right-sided Fisher’s exact test of the hypothesis that the focal class is neutral (i.e., *w*\*=1); see Methods for details.

| Type | Context | All Wild Isolates | PB306 only | PB306 MA | N2 MA | All MA | $w^*_{All} / w^*_{PB306}$ | $p_{All} / p_{PB306}$ |
| --- | --- | --- | --- | --- | --- | --- | --- | --- |
| SV | Exon | 2866 | 630 | 4 | 6 | 10 | **0.14 / 0.17** | <0.0001 / <0.0001 |
|  | Intron | 11818 | 3031 | 5 | 14 | 19 | **0.31 / 0.44** | <0.0001 / <0.002 |
|  | Intergenic | 11292 | 2470 | 6 | 5 | 11 | **0.52 / 0.62** | <0.04 / >0.09 |
| SNV | Splice Variant | 2840 | 515 | 2 | 1 | 3 | 0.48 / 0.48 | >0.17 / >0.17 |
|  | Disruptive Start/Stop Codon | 1830 | 313 | 2 | 2 | 4 | **0.23 / 0.22** | <0.02 / <0.02 |
|  | Missense Variant | 78191 | 14202 | 11 | 30 | 41 | 0.96 / 0.96 | >0.42 / >0.42 |
|  | Intergenic Region | 243014 | 42215 | 53 | 88 | 141 | Neutral | - |
|  | Intron Variant | 274542 | 50937 | 52 | 94 | 146 | Neutral | - |
|  | Synonymous Variant | 82326 | 15411 | 6 | 8 | 14 | Neutral | - |
| Indel | Frameshift Variant | 8446 | 1339 | 4 | 5 | 9 | **0.47 / 0.41** | <0.04 / <0.02 |
|  | In-frame Indel | 2905 | 463 | 0 | 4 | 4 | 0.36 / **0.32** | >0.06 / <0.05 |
|  | Intergenic Region | 74279 | 13577 | 20 | 26 | 46 | 0.81 / 0.82 | >0.10 / >0.12 |
|  | Intron Variant | 100183 | 19053 | 32 | 45 | 77 | **0.65 / 0.69** | <0.001 / <0.003 |

#### (b) False negatives

In contrast to identification of false positives, which both can and must be done by brute force, false negatives can be inferred using bioinformatics (Sedlazeck et al. 2018b). We estimated the false negative rate by (*i*) simulating SVs (“pseudo-variants”) in the reference genome and (*ii*) attempting to identify (“recall”) the pseudo-variant in the sequenced sample (see Methods for details of the simulation procedure). For example, a pseudo-deletion introduced in the reference genome should appear as a variant pseudo-insertion in the sequenced sample. We measured recall in both the MA lines and wild isolates, with the latter providing information about the extent to which reference bias impacts false negative rates.

Recall rates differ for different classes of SV (Figure 2, Supplementary Table 4). The average recall rate is high (∼90%) for deletions up to 10 kb. The recall rate of 100 bp insertions is also roughly 90% but declines as length increases. Recall rates were lower in wild isolates compared to the N2 strain, presumably because the *C. elegans* reference genome is N2; the difference is greater in the highly divergent QX1211 strain (Figure 2, Table 2). As a result, slightly more (∼4%) de novo SVs were missed in the PB306 MA lines than in the N2 MA lines. For reasons that are unclear, we miss almost all small inversions (100-bp), whereas we recall roughly a quarter of 100-kb inversions.

**Figure 2.**
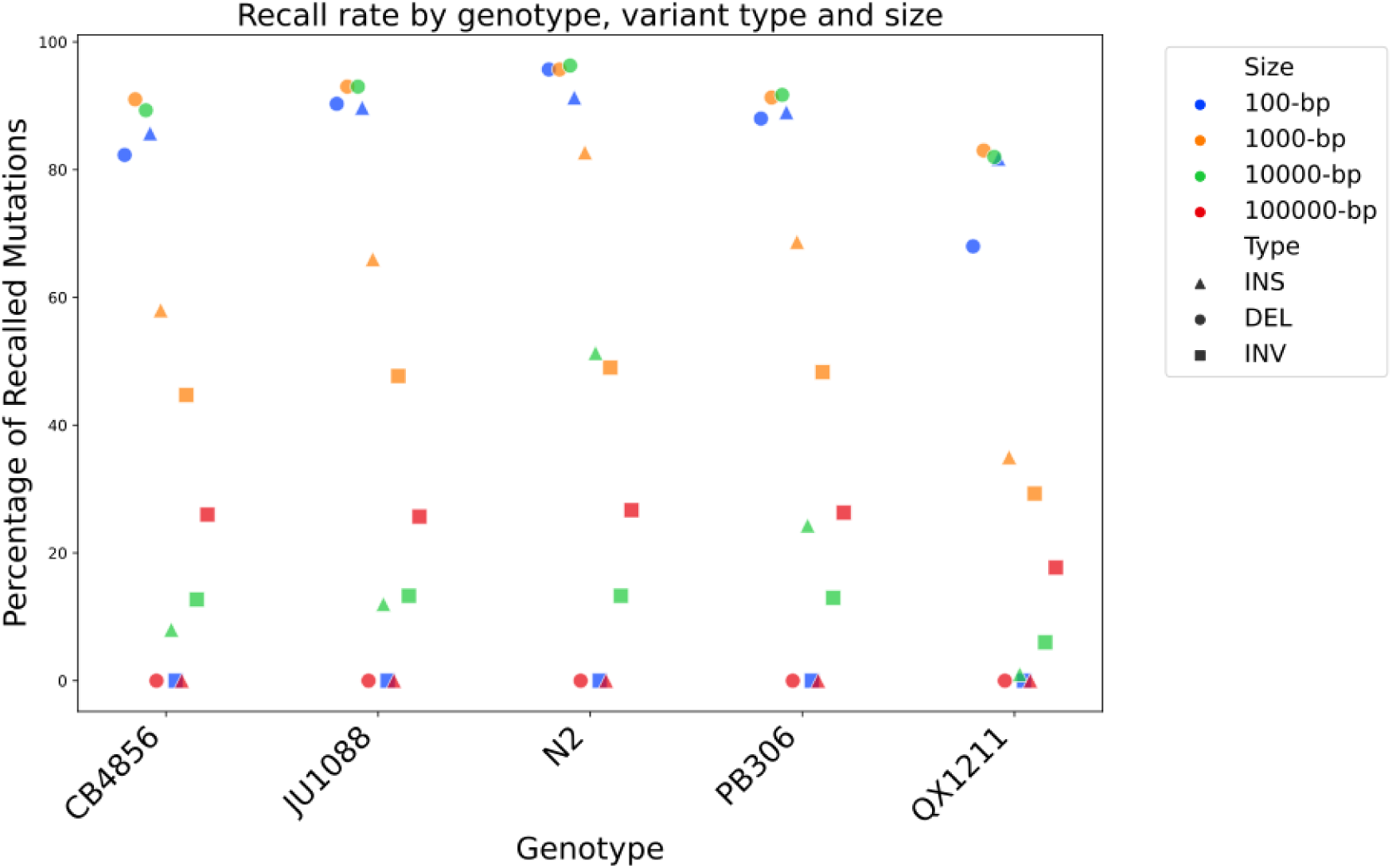
Recall rates by genotype, SV type, and SV size. N2 has the highest recall rate across SV types and sizes. Small deletions (100 bp) have the highest recall rates, comparable to 1-kb and 10-kb deletions. Recall rates of insertions drop sharply beyond 100 bp. Recall rate decreases with increasing SV size and genetic divergence from N2, a trend also observed in PB306-derived MA lines (not shown). Inversions have the lowest recall rates. QX1211, the most divergent genotype from N2, is the most challenging for SV detection. Minor horizontal jitter was added to the x-axis to improve the visual separation of overlapping points.

### SV mutation spectrum

Frequencies of observed SVs are listed by type and average size in Table 2. The total number of bases affected is approximately 16 kb in the four N2-derived lines, with a net loss of 1.5 kb across four lines (7 insertions and 18 deletions). For the two PB306-derived lines, approximately 15 kb of bases were affected in total, with a net gain of 7 kb (5 insertions and 10 deletions). Deletions outnumber insertions 2:1, the same bias observed among smaller indels in a larger set of N2 and PB306 MA lines (Rajaei et al. 2021). Among wild isolates, the number of insertions and deletions are nearly equal, with large insertions outnumbering large deletions by roughly 10% among SVs, and insertions and deletions being nearly equal among smaller indels.

Although our variant caller (PBSV) reports all SVs as insertions or deletions (or inversions, although we did not find any), several of these events represent tandem repeat changes or potentially translocation events. A slight majority of spontaneous SVs (22/40) result from simple repeat expansions or deletions; the average size of the repeat family across 22 independent events is 93 bp. We annotate these cases manually in Supplementary Tables 2 and 3, based on sequence context and alignment features. For example, translocations are typically flagged when inserted sequences show high similarity to distant genomic loci, and tandem repeat changes are identified when insertions or deletions occur within repetitive motifs. These classifications are conservative, as formal proof of translocation or repeat structure would require defining thresholds for sequence identity and repeat purity.

Spontaneous SV mutations are marginally larger in size compared to segregating SVs identified in wild isolates (mean SV size: MA=808 bp, WI=602 bp; median SV size: MA=215 bp, WI=125 bp; Kolmogorov-Smirnov test, p<0.1). This difference is primarily driven by the size of insertions (mean insertion size: MA=1583 bp, WI=799 bp; median insertion size: MA=441 bp, WI=162; K-S test, p<0.05), whereas deletions show no significant difference in size (mean deletion size: MA=475 bp, WI=375 bp; median deletion size: MA=152 bp, WI=105 bp; K-S test, p>0.37). Although the difference in deletion sizes is not statistically significant, we observe a heavier tail in the size distribution of spontaneous deletions, similar to that of spontaneous insertions. This distribution exhibits a multimodal pattern, indicative of larger deletions in the MA lines (Supplementary Figures S1-S3).

Four of the 40 SVs are associated with transposon-related events. In the PB306 MA line 445, a CER1 transposon insertion was identified on chromosome 1, resulting in an 8.8-kb insertion within a short 80-bp intron of the *smg-1* gene. The inserted sequence contains 3’ end of the *plg-1* gene and the full CER1 transposon sequence. Similarly, in the N2 MA line 566, an RTE transposon insertion was observed, leading to a 3.2-kb insertion at the 5’ end of the *alg-1* gene. 3/4 TE events are >3 Kb, whereas the largest non-TE event is ∼2700 bp. The *C. elegans* genome is replete with TEs (Larrichia et al. 2017; Bush et al. 2025), so the greater average size of spontaneous SVs relative to standing variants is unlikely due to an atypically high degree of TE activity in our MA lines. On the contrary, the size discrepancy implies that most TEs are effectively removed by purifying selection.

### Distribution of SVs across the genome

The distribution of spontaneous SV mutations among exons, introns, and intergenic regions does not deviate significantly from expectations based on genomic composition (10 SVs in exon, which constitute ∼32% of the genome; 19 SVs in intron, ∼35%; 11 SVs in intergenic regions, ∼33%; 3x2 goodness-of-fit test, χ²=2.76, df=2, N=40 SVs, p>0.25). Seven of the 40 SVs are located within hyperdivergent genomic regions, consistent with their frequency in the *C. elegans* genome (∼20%; Lee et al. 2021).

In the wild isolates, the distribution of segregating SVs across the genome mirrors the SNV distribution – higher in chromosome arms and lower in the centers (Figure 3); see Figure 3 of Andersen et al. (2012) for the distribution of SNVs along *C. elegans* chromosomes. The recombination rate is higher in chromosomal arms (of autosomes) than centers in *C. elegans* (Rockman and Kruglyak 2009). The greater genetic diversity in chromosomal arms is consistent with reduced Hill-Robertson interference in the arms compared to the chromosome centers. We do not see this pattern on the X chromosome, which is also consistent with expectation, since the recombination rate is nearly uniform on the X.

**Figure 3.**
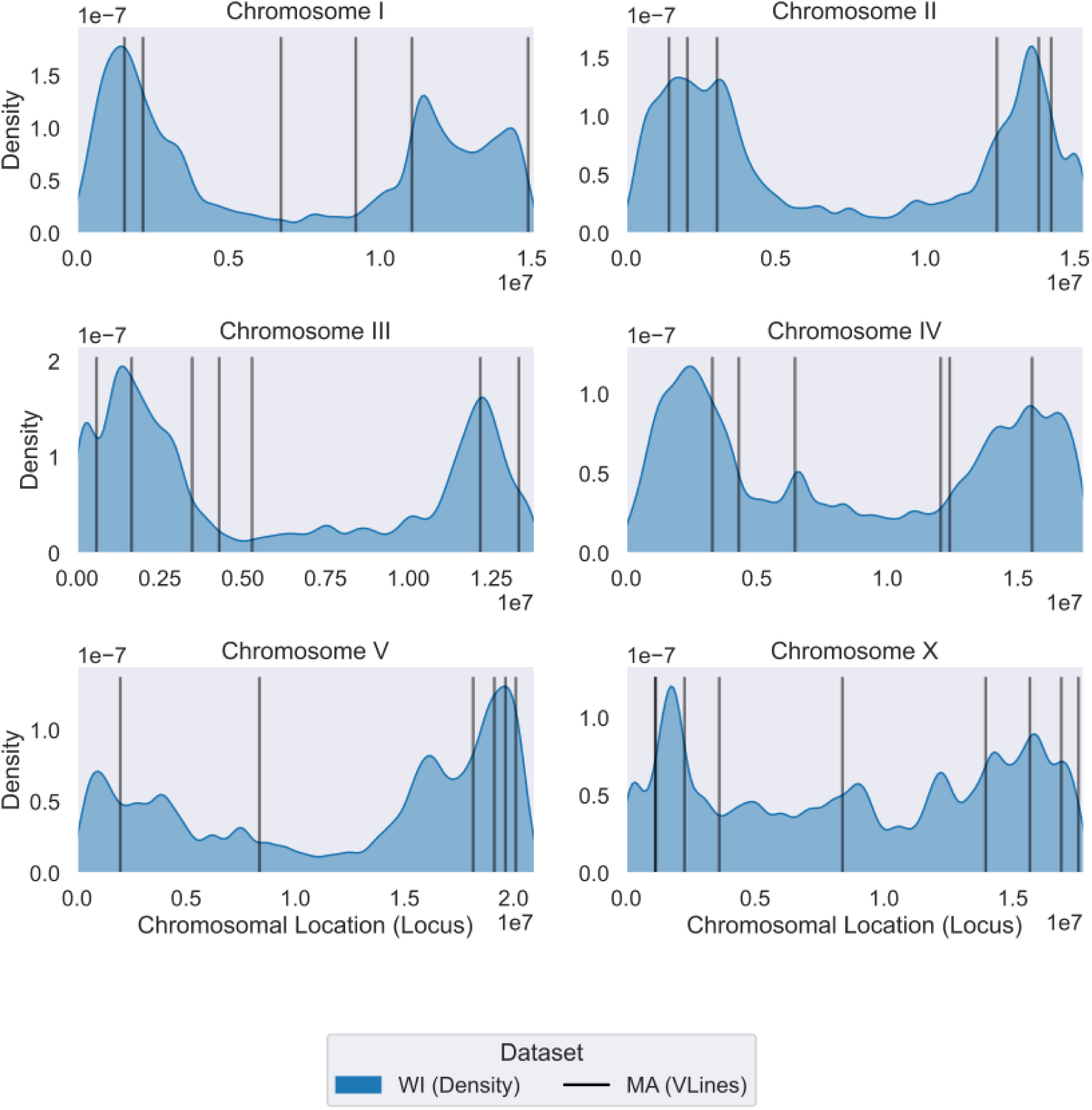
Distribution of SVs within and among chromosomes in wild isolates (density shown in blue) and MA lines (vertical black lines). The overall distributions between these two groups are not significantly different (goodness-of-fit test; See text for details).

It is possible, however, that the chromosomal distribution of genetic diversity is not solely due to variation in the efficacy of purifying selection; it may be that the SV mutation rate is higher in the arms. Indeed, we have previously shown that the base-substitution mutation rate is higher in the arms of the chromosomes than the center in *C. elegans* (Saxena et al. 2019). This view is supported by the fact that low complexity DNA is overrepresented in the arms of *C. elegans* chromosomes (see Supplementary Figures S4 and S5). Even with the small sample size of this study, we observe significantly more spontaneous SVs in the arms of the chromosomes than in the centers (26 SVs in arms which comprise 46% of the genome vs. 10 SVs in the center which comprise 47% of the genome, excluding tips; χ²=7.5, df=1, N=36, p<0.007). In contrast, the distribution of SVs in wild isolates did not differ significantly from the distribution of SVs in MA lines (20,174 SVs in arms, and 6,723 SVs in center, χ²=0.14, df=1, N=26,993, p>0.7). Including chromosomal tips did not alter these conclusions. Genomic boundaries for chromosomal arms and centers follow Table 1 of Rockman et al. (2009). It may be that the distribution of SVs across the genome can be explained by a combination of mutation (more in the arms than the centers) and selection (stronger Hill-Robertson interference in the centers).

### Fitness effects of SVs

To infer the fitness effects of SVs, we compared the spectrum of spontaneous SVs to that of wild isolates (Table 3). Spontaneous SVs are marginally larger than segregating SVs, consistent with the expectation that larger SVs are more strongly deleterious and thus preferably removed by natural selection.

For a more fine-scaled assessment of fitness effects, we compared relative diversity (*w\**; see Equation 1 in the Methods) of different categories of variants between MA lines and wild isolates (Table 3). SNVs in introns, intergenic regions, and synonymous SNVs in exons were pooled into an (assumed) neutral class of variants for this analysis. Unsurprisingly,

SVs in exons are under strong purifying selection; *w\** in exons is approximately 14% of the neutral expectation (p<0.0001, 2x2 Fisher’s exact test). *w\** in introns is ∼30% of the neutral expectation (p<0.0001, 2x2 Fisher’s exact test). Perhaps more surprisingly, *w\** of SVs in intergenic regions is approximately half of the neutral expectation (*w\**=0.52, p<0.04; 2x2 Fisher’s exact test), suggesting strong purifying selection against SVs even in intergenic regions. This observation is, to our knowledge, the first of its kind and warrants further investigation into the functional impacts of intergenic SVs, which may have significant implications for human genetics research. We discuss the implications of this result to the concept of ‘junk DNA’ later.

The preceding calculations are potentially biased, because the MA lines (4 N2, 2 PB306) are more similar to the (N2) reference genome than are the wild isolates, and the failure to recall rate (FTR) increases with divergence from the reference genome (Figure 2). To investigate the possible role of reference bias, we repeated the calculations including only standing variants in the PB306 genome. PB306 is relatively close to N2 and shares 2/6 MA lines, so the effects of reference bias should be less in PB306 than in the average of the wild isolates. The results are qualitatively similar (Table 3), although the values of *w\** are slightly greater (i.e., closer to the neutral expectation) in PB306 than in the full set of standing variants, and the relative paucity of SVs in intergenic regions no longer meets the criterion of statistical significance (*w\**=0.62, p>0.09)

## Discussion

### Rate, Spectrum, and Selective Effects of Structural Variants

The two primary findings of this study are (1) SVs constitute at least 7.5% of spontaneous mutations in the *C. elegans* genome, and (2) SVs are efficiently removed by selection, apparently even from non-coding regions of the genome. But also (3), it is evident that accurately calling de novo SVs is a difficult task that requires a lot of ad hoc curation, even on such friendly terrain as long sequence reads at high coverage of a relatively small, extremely well-characterized, nearly completely homozygous genome.

Robust estimates (i.e., from long-read sequencing) of spontaneous SV mutation rates are now available from *E. coli* (Zhou et al. 2023), the unicellular green alga *Chlamydomonas reinhardtii* (Lopez-Cortegano et al. 2023), mouse (López-Cortegano et al. 2025) and humans (Noyes et al. 2022; Porubsky et al. 2025). *E. coli* and humans span approximately four orders of magnitude of effective population size (Ne ≈ 10^4^ for humans, ≈ 10^8^ for E. coli; Sung et al. 2012), with the accompanying difference in the effectiveness of selection (the “drift barrier”; *s*>1/*N_e_*). The per-genome, per-generation rate of single nucleotide (SNV) mutations also varies by roughly four orders of magnitude between *E. coli* (∼10^-3^/genome/generation) and humans (∼30/haploid genome/generation). SVs constitute approximately 30% of spontaneous mutations in *E. coli* (Zhou et al. 2023) and 6-12% in *Chlamydomonas* (Lopez-Cortegano et al. 2023), but only about 2-3% in humans (Porubsky et al. 2025) and mouse (López-Cortegano et al. 2025). The same calculations for *C. elegans* reveal a per-genome, per-generation SNV mutation rate of ∼0.25/genome/generation and an SV mutation rate of ∼0.03/genome/generation, which corresponds to ∼7.5% of all spontaneous mutations (∼6% using the >50bp threshold) observed across the six MA lines in this study (Table 3; Supplementary Tables 2 and 3). The data are obviously sparse, but the trend is evidently toward a positive relationship between *N_e_* and the fraction of spontaneous mutations that are SVs. Given that SVs are surely more deleterious on average than SNVs, the apparent higher proportion of SV mutations in taxa with more stringent selection is intriguing and seems counterintuitive.

Comparison of the relative diversity (*w\**) between classes of variants provides a heuristic test of the relative strengths of selection. For neutral variants, E[*w\**] = 1 (Equation 1 in the Methods); the smaller *w\** is, the more effectively are variants removed from the population by selection. Comparison of relative diversity (w*) across genomic features (Table 3) shows that structural variants (SVs) exhibit substantially reduced diversity in exons compared to missense SNVs, consistent with stronger purifying selection against exon-disrupting SVs. This is not a surprise; a large indel in an exon is undoubtedly more deleterious on average than an amino acid replacement. Selection also effectively removes SVs from introns. Somewhat less obviously, selection apparently is also effective at removing SVs from intergenic regions. *N_e_* of *C. elegans* is on the order of 10^4^ (Teterina et al. 2023), which implies that many intergenic SVs experience selection *s*>10^-4^.

### Limitations in Estimating the Contribution of Structural Variants to Fitness Decline

Over the course of an average of 238 generations of mutation accumulation (MA), the six MA lines in this study exhibited an average decline in absolute fitness of ∼15% (Baer et al. 2006). Structural variants (SVs) constitute only ∼7.5% of all detected variants. However, accurately quantifying their contribution to fitness decline is challenging without detailed knowledge of the distribution of fitness effects (DFE), which is likely influenced by factors such as variant size and genomic context.

For instance, SVs occurring in exons represent a mutation class with particularly low diversity in natural populations. After adjusting for differences in mutation rates across variant classes, SV diversity in exons is approximately 14% of the neutral expectation. Even that is an underestimate, however, given that the standing variants potentially include relatively new deleterious mutations that have persisted long enough to be observed but are on their way to being selected out of the population (i.e. mutation-selection balance).

In contrast, small indels in introns, the most abundant non-neutral mutation class in our dataset, exhibit diversity at ∼65% of the neutral expectation. Despite this, the MA lines contain approximately eight times as many small indels in introns than SVs in exons. While it is difficult to estimate relative selection coefficients (*s*) for these two classes without a well-characterized DFE, their relative abundance suggests that small indels in introns could contribute substantially to fitness decline, even if their average effect is weaker than that of SVs in exons. Ignoring variant frequency in favor of a focus on average effects risks underestimating the evolutionary consequences of more frequent but potentially less deleterious mutations, such as small indels.

### Effect of False Negatives in Inference of SV Spectrum and Estimates of Selection

Interpretation of the SV spectrum must also be considered in light of the detection limits of our variant-calling pipeline. Our workflow reliably detects variants between ∼30 bp and 2 kb, with recall rates >80% for 1 kb deletions and ∼60–80% for 1 kb insertions (Figure 2; Supplementary Figures S1–S3). These recall rates are similar in both N2 and the wild isolates, with the exception of the QX1211 line, which reduces concern about reference-strain bias within this size range. Detection declines sharply for insertions >10 kb, though such large events account for only a tiny fraction of all calls.

Because the overwhelming majority of variants in our dataset fall within the 50 bp–2 kb range (∼90% of spontaneous variants and ∼95% of segregating SVs), our conclusions about the SV spectrum and the strength of selection necessarily pertain to this size class. Because we did not detect many variants much larger than ∼2 kb, we avoid making inferences about mutation or selection acting on larger SVs for which our detection power is limited.

### Structural Variants in Low-Complexity Mutational Hotspots

A significant proportion (∼22.5%; 9 out of 40) of structural variants (SVs) identified in mutation accumulation (MA) lines occur in low-complexity mutational hotspots. Previously, Konrad et al. (2018) reported a similar estimate (∼31%) for duplications and a much higher estimate (∼81%) for deletions occurring in loci with pre-existing CNVs in the ancestor, based on short read sequencing of N2-dervied MA lines in *C. elegans*. Among the SV calls in MA lines, the majority (22 out of 40) are simple repeat expansions or deletions. These annotations were determined through manual inspection of variants using the JBrowse genome browser as well as IGV rather than automated annotation, which, while scalable, may introduce noise. The repeating units range in size from as small as dinucleotide repeats (e.g., AT, CA, CT) to larger structural events, such as the deletion of a single copy within a four-copy 360-mer repeat.

An unexpected observation is that 9 of the 22 SVs identified in repeat-rich regions required a ‘manual correction’ due to a common misclassification in the workflow – where the MA variant call occurs on a locus with a pre-existing variant in the ancestor. This suggests that these loci represent mutational hotspots. Notably, the variant caller PBSV did not correctly handle these cases, even in joint genotyping mode. Supplementary Table S2 and S3 provides annotations for SV calls in MA lines, detailing the necessary corrections following manual verification using IGV and genome browser inspection.

For example, a variant call on chromosome I of PB306-derived MA line MA445 (I: 1567493), illustrated in Figure 4a with a pedigree and 4b with an IGV snapshot highlights this issue. In the PB306 ancestor, this region contains a 3.1 kb insertion composed of perfectly repeating copies of a 173-mer interspersed with additional sequence. Some of this additional sequence contains similar repeating motifs, but of smaller units, and precise annotation is difficult due to sequencing errors. The sequence of this insertion, along with its repeating unit, is provided in Supplementary Sequence File S1 in FASTA format. We show the alignment of raw PacBio reads in Supplementary Figure S6 and show the repeating copies of 173-mer highlighted in text in Supplementary Figure S7. The same 3.1 kb insertion is also present in the PB306-derived MA line MA459, consistent with the expectation that a variant present in the ancestor is inherited by its descendant. This insertion is absent in the N2 ancestor, indicating that it is polymorphic in *C. elegans*. Interestingly, in MA445, this region harbors a 1.9 kb insertion, effectively representing a 1.2 kb deletion relative to the PB306 ancestor. However, variant calling software failed to annotate this as a deletion event, necessitating extensive manual validation of both raw and assembled reads. Because some repeat copies are perfectly conserved while others exhibit minor variations, the identification of this type of event remains beyond the resolution of current variant calling algorithms.

**Figure 4.**
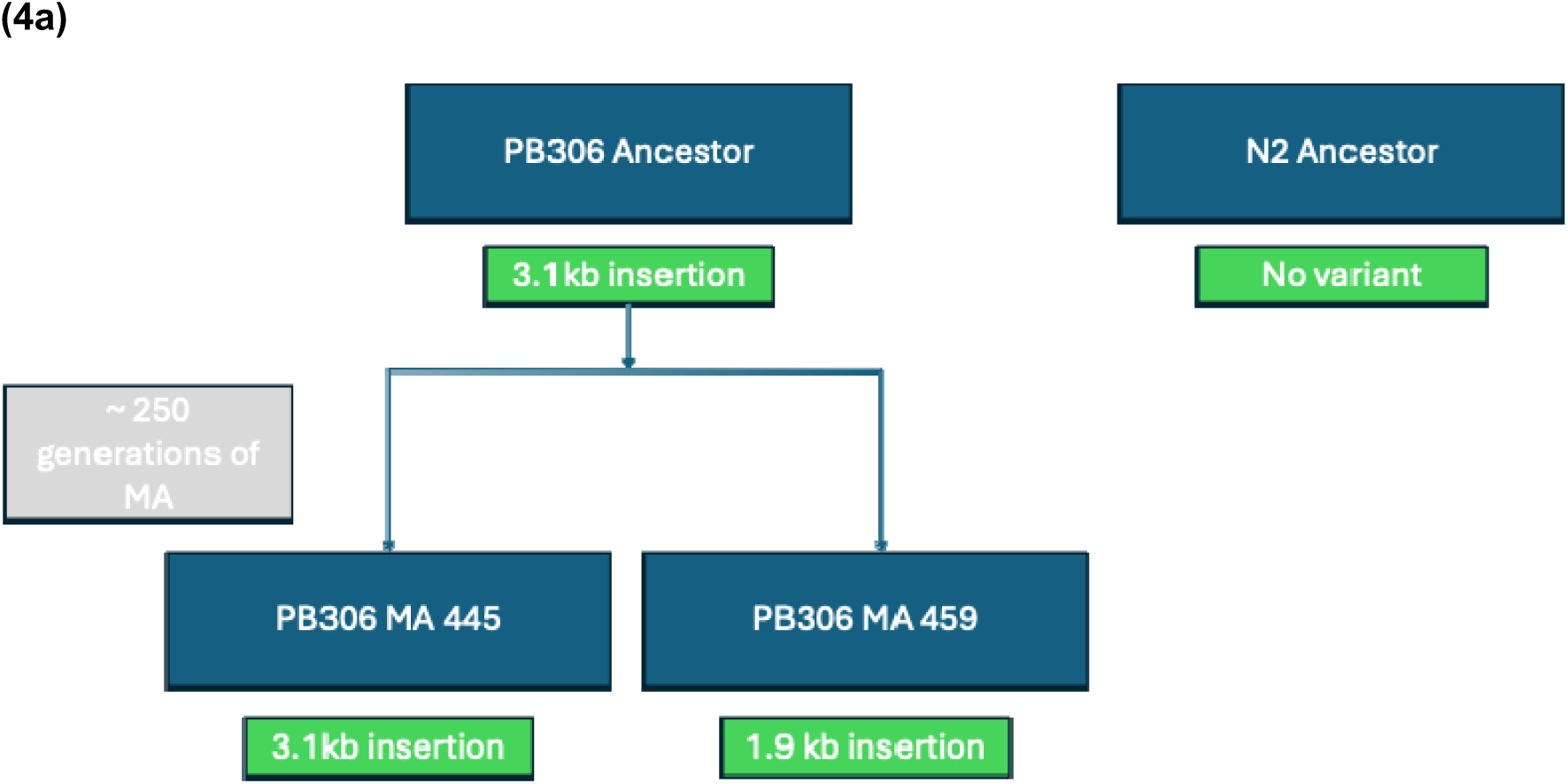
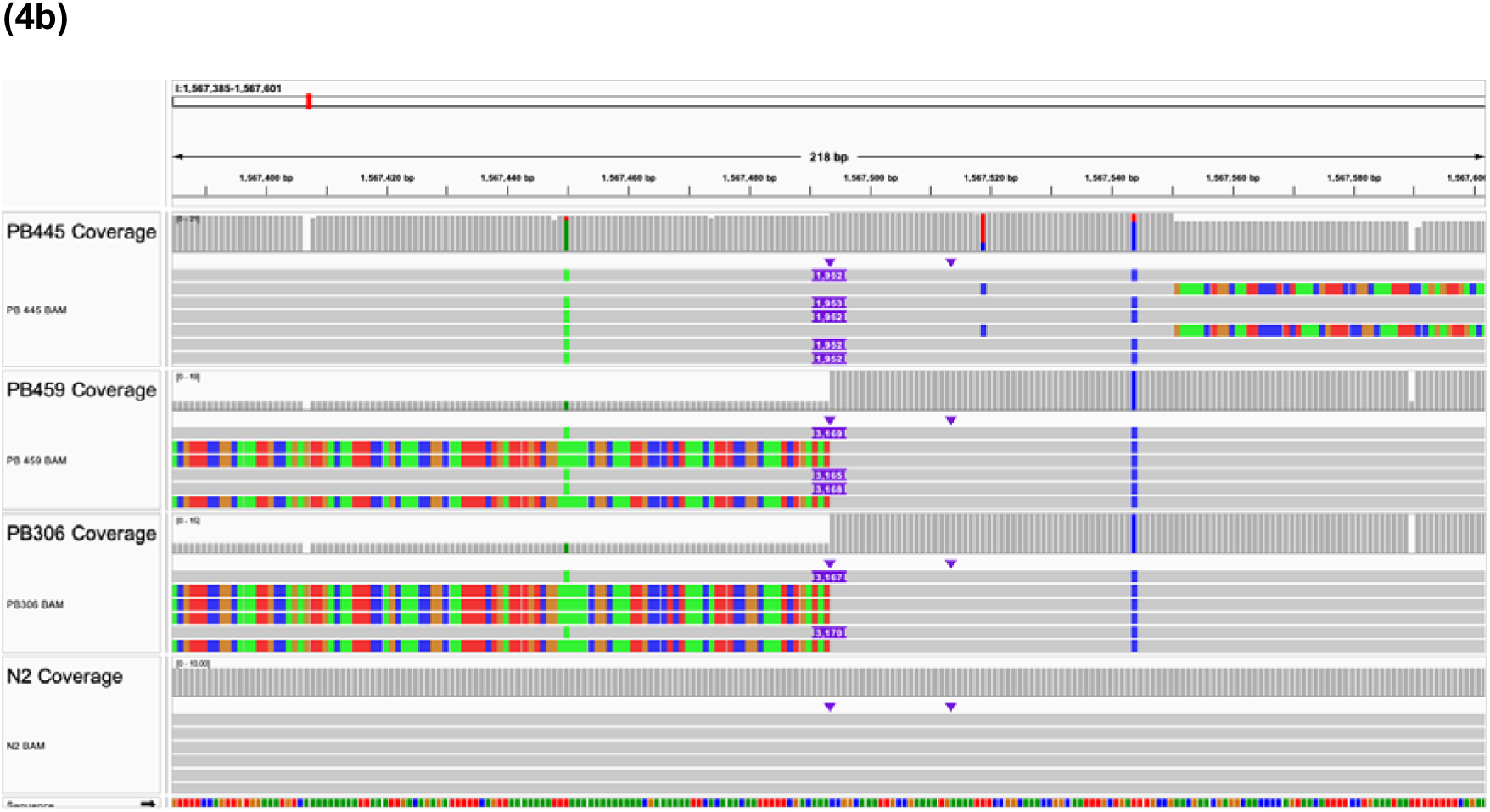
(Top) A schematic representation of the variants identified at the locus (I: 1567493) in the pedigree of MA lines. PBSV initially classified the 1.9 kb insertion and the 3.1 kb insertion as two independent mutations. However, a deeper analysis—accounting for sequencing errors—reveals that a 1.2 kb segment was deleted within the inserted sequence in the ancestor, leading to the observed 1.9 kb insertion. (Bottom) IGV snapshot of the 1.2 kb deletion in PB306-derived MA 445. The IGV snapshot shows pseudo-read alignments, which are assembled contigs representing longer sequences. Some pseudo-reads fail to fully capture the inserted segment and appear as soft-clipped sequences. If the inserted segment is not entirely covered within the pseudo-read, it is clipped from the alignment, appearing as base calls (colored bars) instead of clean (gray bars), fully aligned sequences. Some insertion sequences are misaligned because of the repetitive nature of the locus; however, all pseudo-reads carry the insertion. Supplementary Figure S6 presents the same analysis using error-prone PacBio raw reads.

The need for manual correction in 9 out of 22 repeat-associated SVs raises an important question: Are certain repeat regions disproportionately prone to structural variation? The *C. elegans* genome contains a vast number of repeat-associated sequences. We observed 122,161 distinct repeat-associated sequences in the WS270 GFF3 file that span roughly 19.5% of the genome (See Methods). Despite this substantial genomic footprint, only a small subset of these regions harbored recurrent SVs in our lines. Our findings suggest that loci already carrying an SV in the ancestor are prone to recurrent SVs, making them significant hotspots for genetic variation. Analogous SV mutational hotspots have been identified in the human genome (Porubsky et al. 2025). Furthermore, these regions are likely underrepresented or misannotated by automated variant calling pipelines. With only 9 such identified events, it is beyond the scope of this study to understand the factors that cause some repeat sequences to be hotspots. Notably, 2 of the 9 manually corrected repeat-associated SVs occur in hyperdivergent regions—an observation worth noting, though we refrain from drawing broader conclusions.

### Estimating the Fraction of the Genome under Evolutionary Constraint

We observe that selection efficiently removes most SVs, regardless of their genomic locations (genic versus intergenic). This finding challenges the conventional view of “junk DNA,” which is often assumed to evolve free from the constraints of purifying selection. Our results suggest that a larger fraction of the genome may be functionally relevant than previously inferred from studies focusing on base substitutions. Unconstrained genomic regions are typically defined by the absence of purifying selection. However, it is possible that some genomic regions can tolerate base substitutions but not SVs. For instance, consider a hypothetical low-complexity region between an enhancer and a promoter (Bateman and Johnson 2022). Base substitutions in such regions may not disrupt gene function, whereas SVs that alter the distance between the enhancer and promoter might, which could in turn negatively impact fitness. Similarly, regulatory sequences flanking genes may be more tolerant of base substitutions than of small indels or larger SVs. In a similar vein, Payer et al. (2017) found that polymorphic *Alu* elements in the human genome were significantly associated with Trait Associated SNPs (TASs) in human GWAS. Larger-scale studies, capable of identifying more SV mutations, could reveal many such regions that appear to evolve freely based on SNP variation but are subject to purifying selection against SVs.

Taken together with recent findings from other organisms, the results reported here have implications for the design and interpretation of studies of complex traits in humans (e.g., GWAS). The most recent estimates of *N_e_* in *C. elegans* – on the order of 10^4^ (Teterina et al. 2023) – are surprisingly close to estimates of *N_e_* in humans (Tenesa et al. 2007), given that 10^8^ *C. elegans* could fit comfortably inside a single human. A recent study with yeast (*S. cerevisiae*) revealed that including SVs in GWAS explained an additional ∼14% of heritable variance in thousands of traits, relative to the variance explained by SNVs alone (Loegler et al. 2025). If even intergenic SVs are under effective selection (*s*>1/*N_e_*) in *C. elegans*, it implies the same may be true for humans, all else equal. Which in turn implies that SVs are likely to explain significant additional heritable variance for human complex traits, but also that those variants are likely to be rare.

## Conclusion

The spontaneous SV mutation rate is inferred to be approximately 7.5% of the SNV/small indel rate, but that is likely an underestimate. SVs are efficiently removed from natural populations, indirectly implying a function (biochemical or otherwise) to a larger fraction of the genome than is typically assumed.

## Material and Methods

### MA Protocol

The MA protocol has been described in detail previously (Baer et al. 2005) and is depicted in Figure 1. The maximum number of MA generations (Gmax) is 250. The method by which we estimated the number of generations of MA for a line (below Gmax) is reported in Saxena et al. (2019).

### DNA extraction, library preparation, and sequencing

Cryopreserved samples of the MA lines were thawed onto 100 mm NGMA agar plates seeded with a lawn of *E. coli* OP50 and grown for 2-3 days, until there were many gravid worms, at which time worms were harvested, “bleached” to remove bacterial and fungal contamination (Stiernagle 2006), and embryos transferred to a new, seeded 100 mm plate. L1 larvae were collected the next day and transferred to seven seeded 100 mm plates and grown until the plates were nearly starved. Worms were washed from the plates, double-rinsed in M9 buffer, and pelleted for DNA extraction. DNA was extracted from each sample with the Quiagen Gentra Puregene kit, following the manufacturer’s instructions. Extracted DNA samples were sent to the University of Maryland genomics core facility for library preparation and sequencing on the Pacific Biosciences Sequel platform.

### Variant Calling Pipeline

We evaluated multiple variant calling strategies, including direct alignment-based approaches and de novo assembly followed by genome-to-genome alignment using Assemblytics (Nattestad and Schatz 2016). Ultimately, we adopted a modified version of the SMRT-SV2 (Audano et al. 2019) pipeline, as it provided a balance between sensitivity and specificity while minimizing false positives. A detailed discussion of our rationale for selecting this approach over others is provided in section 1 of the Supplementary Discussion.

The SMRT-SV2 pipeline processes error-prone PacBio subreads by aligning them to the reference genome using blasr (Chaisson and Tesler 2012). These alignments are then segmented into 60-kb tiles, which are assembled using Canu (Koren et al. 2017). This localized assembly approach reduces interference from reads originating from other genomic regions. Importantly, the assembled contigs effectively “clean” the PacBio subreads, leading to improved downstream alignments. While SMRT-SV2 automatically re-aligns these contigs using blasr, we opted to replace blasr with minimap2 (Li 2018), which provided superior alignment performance for our dataset.

SMRT-SV2 includes its own variant caller. However, it led to an unacceptably high false positive rate for de novo mutations. To improve accuracy, we benchmarked several alternative structural variant (SV) callers, including Sniffles (Sedlazeck et al. 2018a) and PBSV (Wenger et al. 2019), ultimately selecting PBSV due to its optimized performance on high-quality PacBio reads. Further details on the performance of SMRT-SV2, Sniffles, and SMRT-SV2 variant calling are provided in section 3 of the Supplementary Discussion.

In cases where a potential variant was detected, SMRT-SV2 produced multiple assemblies—referred to as “pseudo-reads”—for the candidate locus. These pseudo-reads were then aligned with minimap2 and variants were called using PBSV. Despite this refinement, we still observed many false positives, albeit a manageable number, further necessitating manual curation. Each candidate variant was inspected in IGV (Robinson et al. 2017) and JBrowse genome browser (Diesh et al. 2023), comparing error-prone raw PacBio subread alignments, pseudo-read alignments, and independent Illumina sequencing data generated for a separate study. Only variants present in the mutation accumulation (MA) line but absent in both the MA ancestor and other MA lines were considered true de novo mutations. Once validated, variants were functionally annotated using genome browser annotations to classify their genomic context (e.g., exon, intron, UTR, repetitive element, hyperdivergent region, potential regulatory elements).

For identifying segregating SVs, we followed the same pipeline with one key difference: instead of an ancestor-descendant comparison, all wild isolates were jointly called using PBSV.

### Estimating Sequencing Base-Call Error rate

Base-call errors arise during PacBio sequencing and manifest as nucleotide-level mismatches between raw subreads and the reference genome. We quantified these errors using Levenshtein distance, which measures the minimal number of single-character edits (insertions, deletions, substitutions) required to transform one sequence (read) into another (aligned segment of reference). Raw subreads were aligned to the reference genome using minimap2 (Li 2018), and Levenshtein distance was computed for each alignment using the ‘NM’ tag in the bam file. We estimated the base-call error rate of pseudo-reads generated after alignment and assembly similarly by aligning the pseudo-read to reference, and parsing the ‘NM’ tag in the bam file. This analysis ignores soft-clipped regions which could contain worse base-call quality on the edges of the reads, hence this analysis provides a conservative estimate of base-call error rate.

### Variant Annotation

Annotation of structural variants (SVs) identified in the mutation accumulation (MA) lines was performed manually. Basic genomic classifications—such as exon, intron, and intergenic regions—were straightforward to assign. Similarly, regulatory annotations, including DNase hypersensitive sites, transcription factor binding sites, and hyperdivergent regions, could be systematically determined using existing datasets. However, annotating repetitive elements within SVs was significantly more time-consuming. Many SVs overlapped repetitive sequences that were not explicitly annotated by RepeatMasker, requiring manual inspection of the variant and read sequences to deduce repeat structure.

In contrast, annotation of SVs in wild isolates was automated using GFF3-based genome annotations. Variant calls were intersected with genomic features using *bedtools* (Quinlan and Hall 2010), allowing for systematic classification. Automated annotation tools such as *snpEff* (Cingolani et al. 2012) were not suitable for SV annotation, as they do not properly account for the full extent of large structural variants. For example, an 8-kb deletion may only be annotated at its breakpoints rather than across its entire span. To ensure accurate functional classification, we assigned the highest-impact annotation observed across the deleted region. If a deletion started upstream of a gene but extended into coding sequences, it was annotated based on the most functionally significant feature it disrupted (e.g., an exon rather than an upstream region).

Short variants (SNPs and small indels) in both MA and wild isolates were annotated using *snpEff* with the following flags: -no-downstream -no-upstream -no-utr. As a result, upstream, downstream, and UTR annotations were instead categorized as intergenic, maintaining consistency with our SV annotation approach. *snpEff* produces a diverse set of annotations, many of which are redundant or overly specific for our purposes. To standardize annotation categories, we applied a systematic simplification approach using a predefined mapping (Supplementary Table 5). This mapping condensed detailed annotation labels into a manageable number of biologically meaningful categories, ensuring consistency across variant types.

### Estimating the relative strength of selection for different categories of mutations

To estimate the relative strength of selection we first defined a neutral class of variants, including intergenic SNVs, intronic SNVs, and synonymous SNVs within exons. We included intronic and intergenic variants in addition to synonymous variants to increase the sample size of MA variants in the neutral class, given the small number of synonymous MA variants in our dataset (n=14). The relative diversity (*w\**) of a focal class of variant (e.g., SVs in exons) was calculated from the equation:

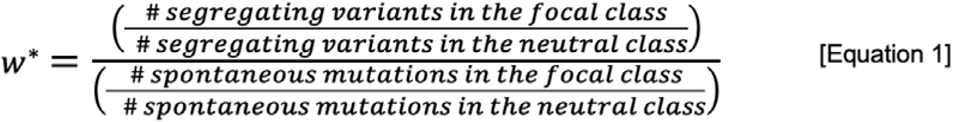

This approach quantifies the standing diversity within a particular class of (focal) variant relative to the diversity in the neutral class, accounting for the mutation rates in both categories. If the focal class of variants is neutral, E(*w*\*) = 1. If the focal class of variant is subject to purifying selection, there will be fewer segregating variants than expected given the mutation rate and *w*\*<1. Statistical significance is assessed by a 2x2 Fisher’s exact test. Counts of variants in each category in the wild isolates and MA lines are given in Table 3.

### Simulation of Reference Genomes and Estimation of False Negative Rates

To evaluate the sensitivity of our variant calling pipeline, we simulated structural variants in the *C. elegans* reference genome (WS270) using the R package *RSVSim* (Bartenhagen and Dugas 2013). For each variant type—insertions, deletions, and inversions—we generated three independent simulated genomes. Within each variant type, we introduced 100 variants of lengths 100 bp, 1,000 bp, and 10,000 bp, as well as 50 variants of 100,000 bp.

Following simulation, we applied our variant calling pipeline to assess recall rates. Introducing structural variants into the reference genome shifts the positions of subsequent variants, making direct locus-based comparisons dependent on tracking the precise order of variant introduction. While this could be addressed if the software explicitly recorded variant placement order, doing so would introduce additional dependencies into the analysis. To avoid this complexity, we instead used a size-based evaluation approach, considering a variant correctly detected if its length fell within predefined tolerance ranges: (98–102) bp, (995–1,005) bp, (9,995–10,005) bp, and (99,995–100,005) bp for the respective size categories. This approach simplified the evaluation while providing a systematic estimate of false negative rates.

### Estimating number of repeat-associated entries in C.elegans genome

To quantify the genomic burden of repetitive DNA, we merged overlapping annotations from the WS270 GFF3 file, including “repeat_region”, “inverted_repeat”, and “tandem_repeat” entries. This yielded 122,161 distinct repeat-associated regions, collectively spanning approximately 19.5 Mb or 19.5% of the *C. elegans* genome.

### Data Analysis

All statistical analyses (e.g., Kolmogorov-Smirnov test, Fisher’s exact test) were performed in Python using SciPy (Virtanen et al. 2020). Data visualization was conducted using Matplotlib (Hunter 2007) and Seaborn (Waskom 2021).

## Supporting information

Supplementary Sequence File S1

## Acknowledgments

We thank Tim Crombie, Joanna Dembek, Lindsay Johnson, and Mike Snyder for assistance in the lab, and Erik Andersen and Annalise Paaby for providing the wild isolate data. Two anonymous reviewers provided insightful comments on a previous version. Support was provided by NIH award GM107227 to CFB.

