## Supplementary Figures for "High rate of mutation and efficient removal by selection of structural variants from natural populations of *Caenorhabditis elegans*"

**Supplementary Figures S1–S3:**
These density plots illustrate the size distribution of structural variants (SVs) in both mutation accumulation (MA) lines and wild isolates. **S1** presents the distribution for all SVs, while **S2** and **S3** specifically depict insertions and deletions, respectively. Notably, all three plots exhibit a **multimodal distribution**, with a smaller secondary peak on the right for **MA-derived SVs**, indicating a distinct size pattern in these variants. For clarity, 117 out of 22307 SVs in the wild isolate set (ranging in size from 10,000 to 90,000 bp) were removed from the plots to allow axis scaling that better displays the overall distribution.

**Supplementary Figures S1**

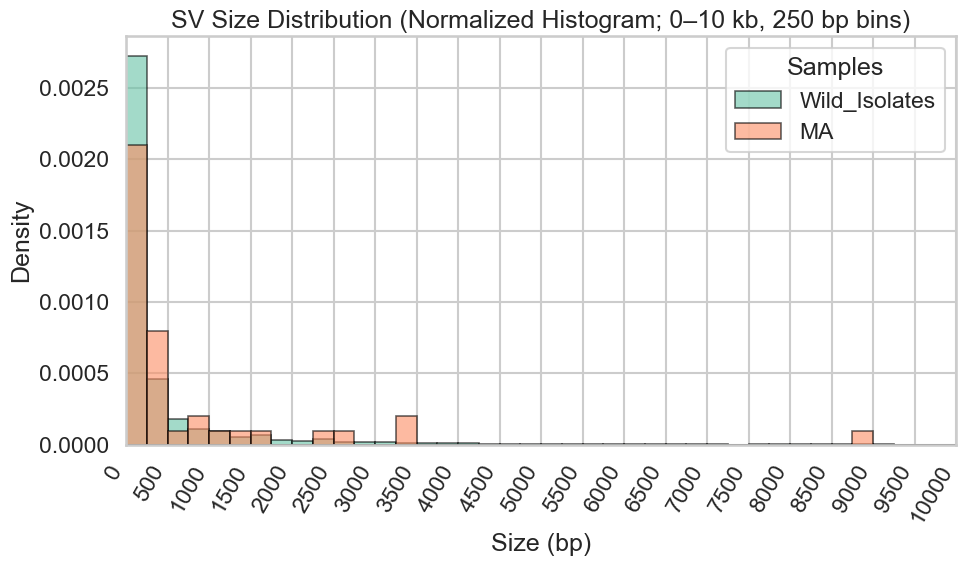

**Supplementary Figures S2**

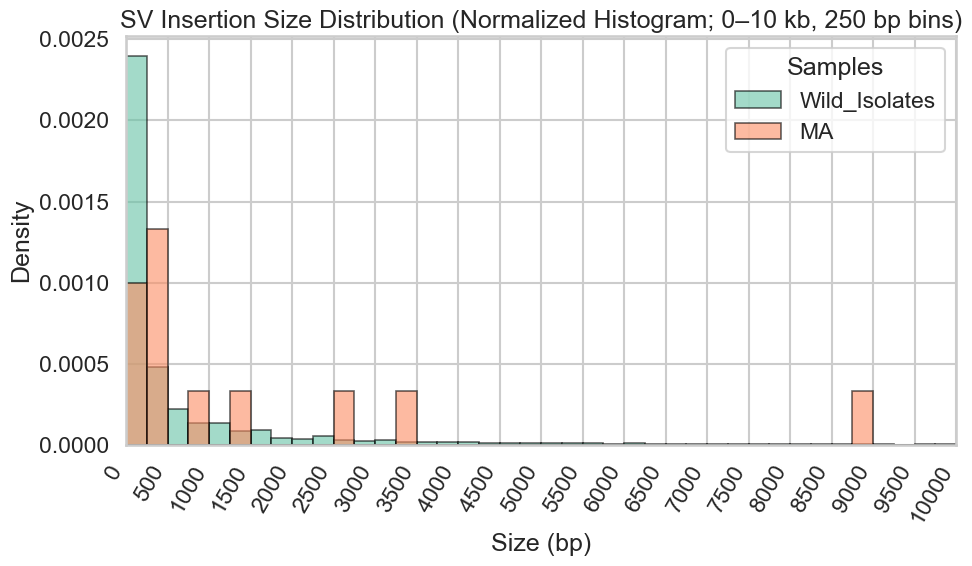

**Supplementary Figures S3**

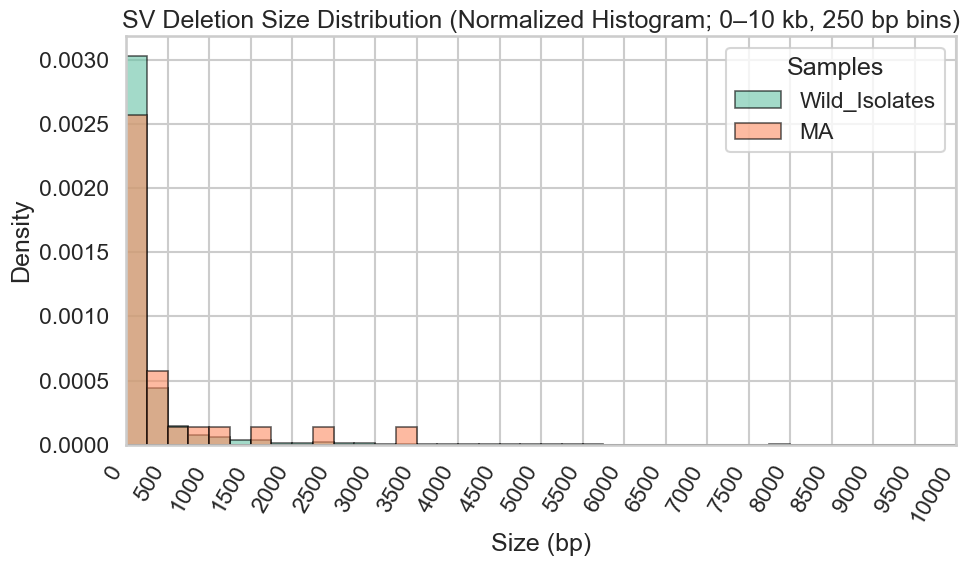

**Supplementary Figures S4–S5:**
These density plots display the chromosomal distribution of **low-complexity DNA** (**S4**) and **transposable elements** (**S5**) across the C. elegans genome (greater in the arms, and lower in the center).

**Supplementary Figures S4**
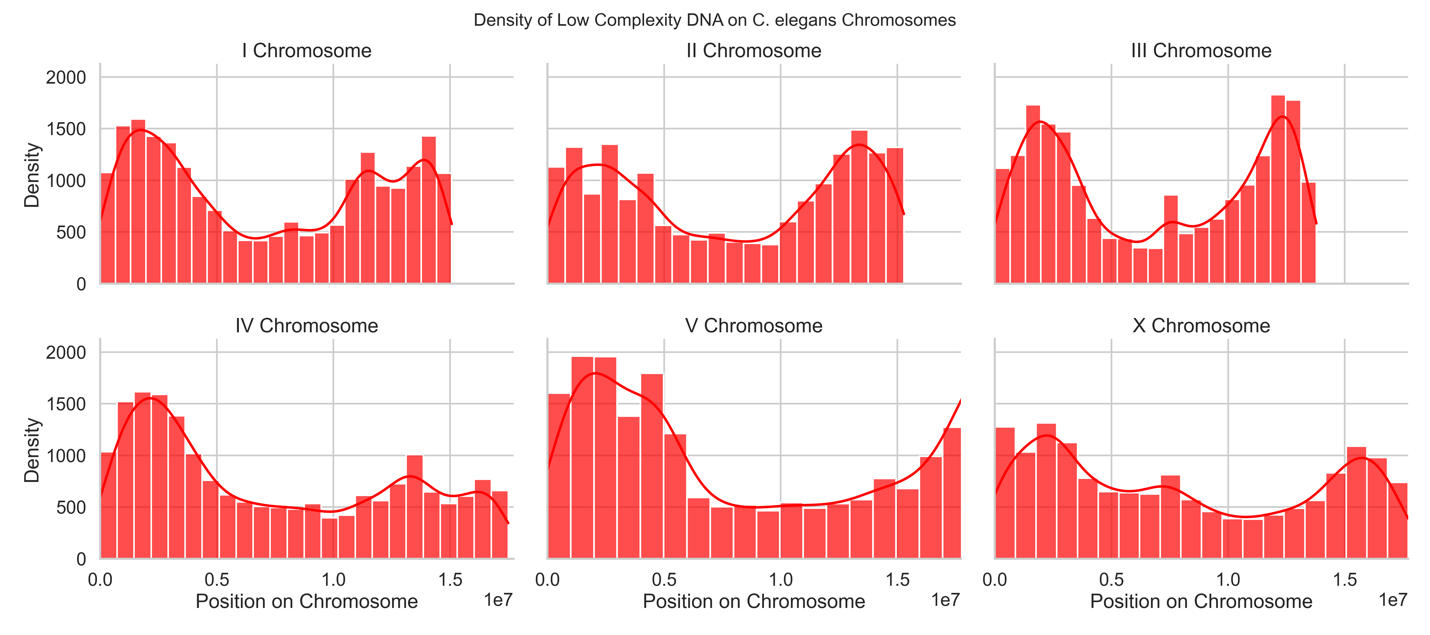

**Supplementary Figures S5**

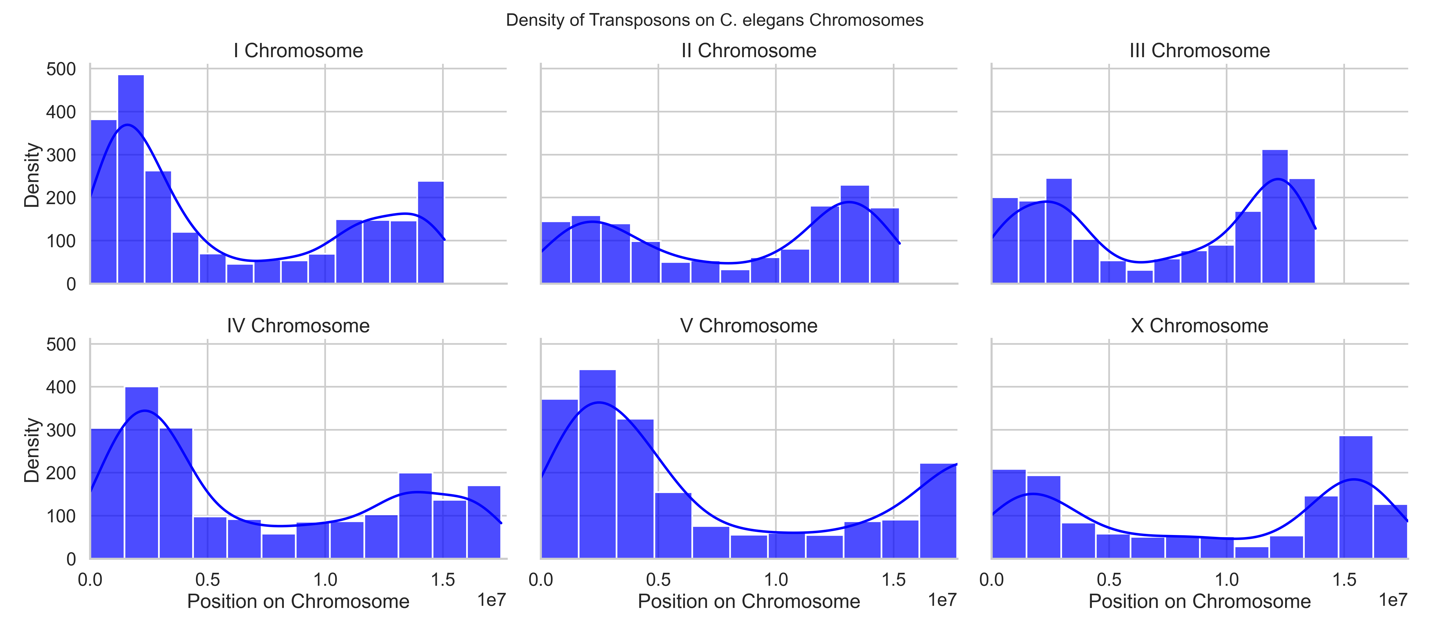

**Supplementary Figure S6:**
An **Integrative Genomics Viewer (IGV) snapshot** showing a **1.2 kb deletion** in the PB306-derived MA line **MA445**. This visualization includes alignment data from **raw PacBio subreads** mapped to the **WS270 genome** for four lines: **MA445, MA459, PB306 ancestor, and N2 ancestor**. A similar figure in the main text presents the same deletion using **pseudo-reads (assembled contigs)** for these lines, offering a complementary view of the variant. Alignments from raw subreads tend to be imperfect because of sequencing errors, with insertions being of slightly different sizes, and have slightly different positions.
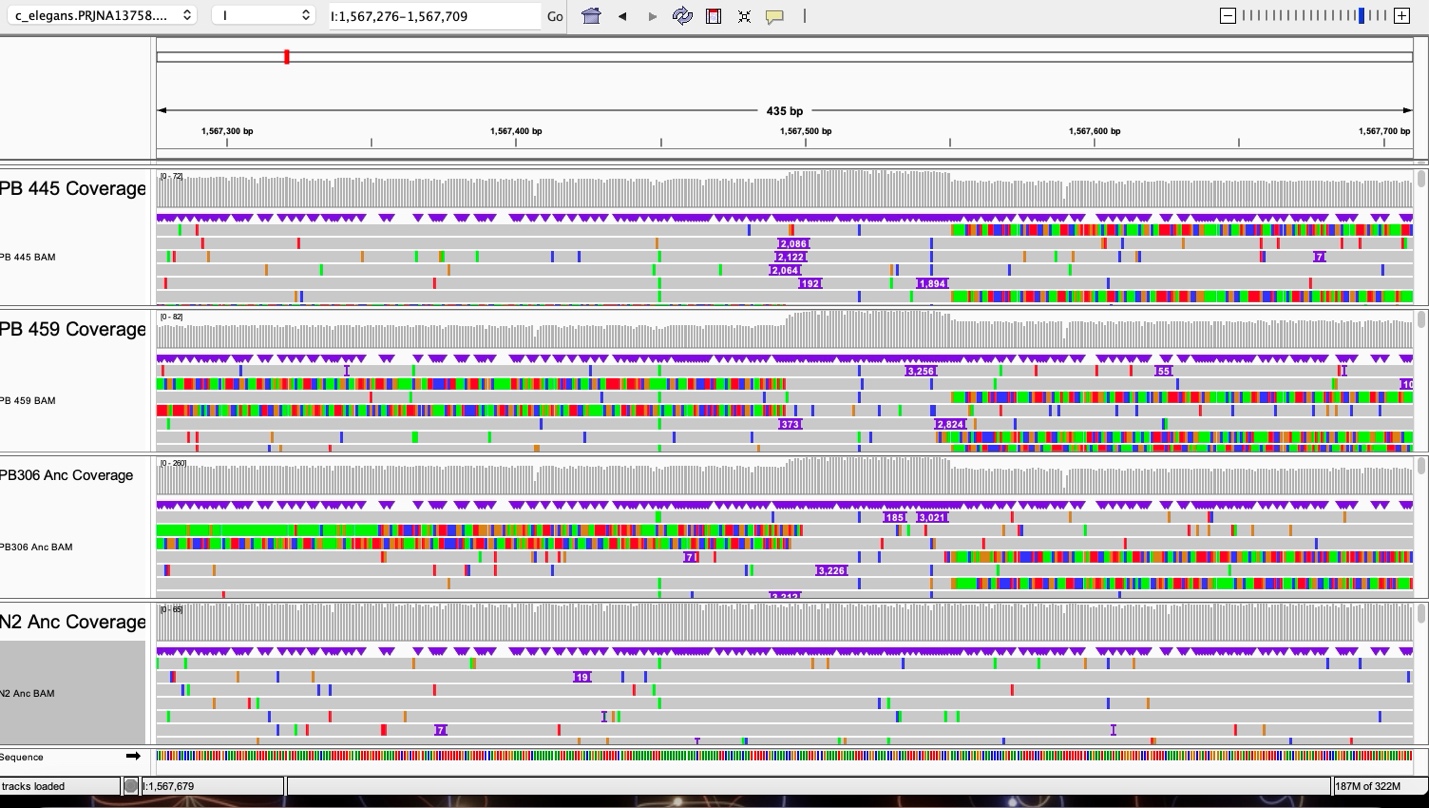

**Supplementary Figure S7:**
A **text-based visualization** of a **173-bp repeating motif** found in two pseudo-reads: one from a **3.1 kb insertion sequence** and another from a **1.9 kb insertion sequence**. This figure highlights the repetitive nature of the sequence, which was previously discussed in relation to structural variation in the dataset. The repeating sequence (bottom) is highlighted and is present in 9 perfect copies in the 3.1kb insertion sequence (top) and 6 perfect copies in the 1.9 kb insertion sequence (middle).
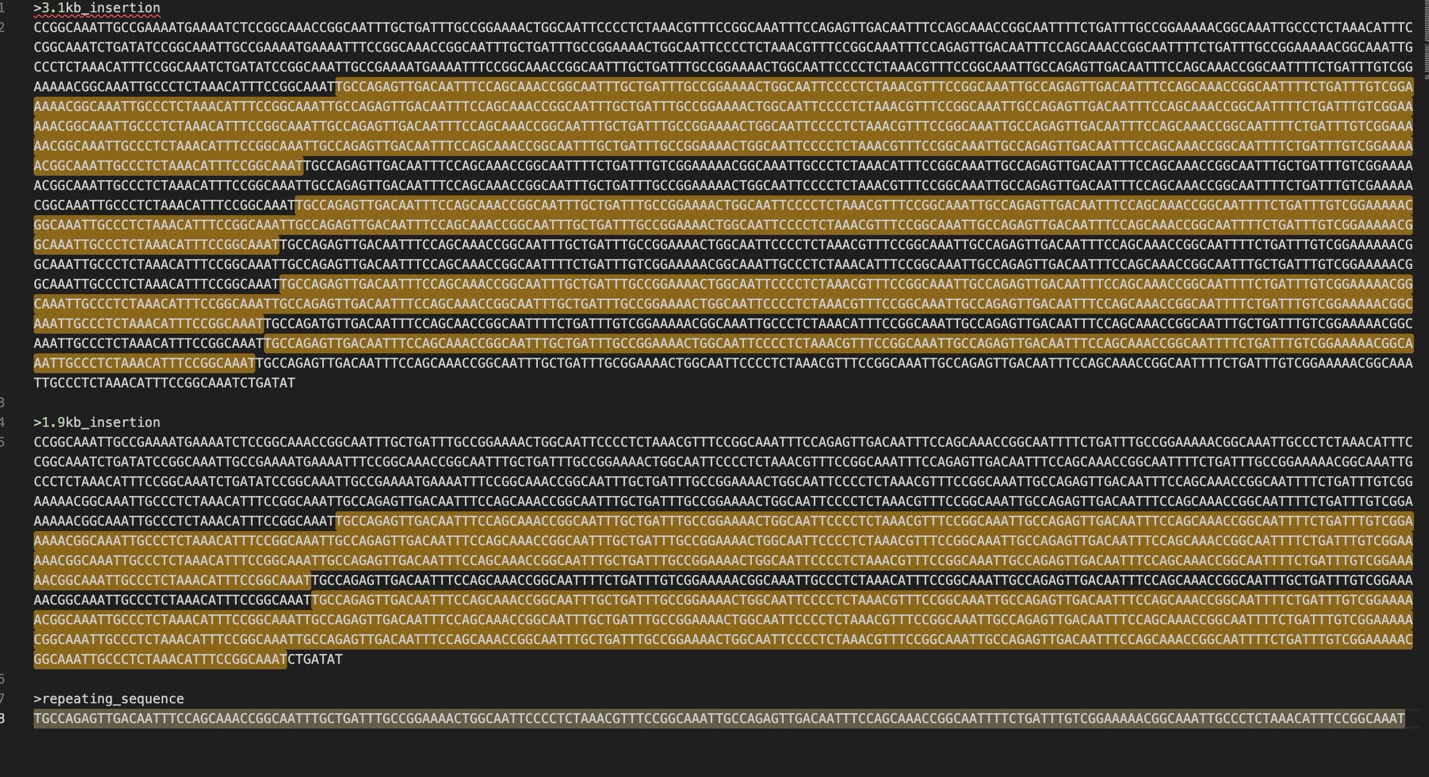
