## Supplementary Material for "High rate of mutation and efficient removal by selection of structural variants from natural populations of *Caenorhabditis elegans*"

**Supplementary Discussion**

**Section 1**

*False Positives and Challenges in Developing an SV Calling Workflow*

Developing an accurate structural variant (SV) calling pipeline requires balancing sensitivity (few false negatives) and specificity (few false positives), particularly when detecting spontaneous mutations. Even the best pipeline will result in some false inferences. However, we expected that if a workflow satisfied certain criteria, the error rates would be low enough to allow reliable SV discovery. Specifically, we expected a well-performing pipeline to meet the following criteria:

1. **Reproducibility in Mutation Accumulation (MA) Lines** – Any variant identified in the MA progenitor (i.e., a difference between the MA progenitor and the reference genome) should also be detected in all of the derived MA lines, provided the region is covered. This is an automatable way of quantifying false positive rates. If a variant appears in the progenitor but is absent in the MA lines, it is likely a false positive in the progenitor. We ignore the possibility of back mutations for this assessment. The power increases with the number of MA lines, because with only a small number of MA lines, the occasional segregating heterozygote in the ancestor may not appear in any of the MA lines. Mischaracterizing ancestral heterozygotes as false positives will bias the inferred false positive rate upward.
2. **Correlation between SV and SNP Diversity** – The nucleotide diversity estimated as the pairwise genetic distance (π) from SVs among wild isolates should correlate with π estimated from single nucleotide polymorphisms (SNPs). A strong correlation would suggest that SV calling captures true population-level variation.
3. **Declining Discovery Rate with Increasing Sample Size** – As more wild isolates are analyzed, the total number of newly discovered SVs should decrease and ultimately reach an asymptote. While this criterion is necessary, it is not sufficient on its own to confirm pipeline accuracy.

Our initial assumption was that a pipeline meeting these three criteria would still contain some false positives and false negatives, but at levels low enough to be tolerable for spontaneous SV detection. However, it turns out that even when all three criteria are met, the pipeline is still inadequate for accurately identifying spontaneous SVs. The fundamental issue is that spontaneous SVs are rare events, such that even a small number of false positives overwhelm the few true positives. For example, if a pipeline generates ten false positives per genome, the impact differs dramatically depending on the genomic context. In wild isolates, which typically harbor around 10,000 true SVs, this corresponds to a negligible 0.1% false positive rate. However, in an MA line that carries only ten new SVs in its genome, the same number of false positives (ten per genome) leads to a false positive rate of 50%. In this scenario, false positives overwhelm the true signal. To re-emphasize an observation we made before, even in our current workflow, MA 517 contained 5 real SVs and 35 false positives, although this is a high outlier.

*Why the First Criterion Failed*

At first glance, variants present in the MA progenitor but absent in the derived MA lines should indicate a false positive (or a residual heterozygote) in the progenitor. However, many of these cases were due to assembly artifacts, which were stochastically called in one sample (ancestor) but not another (MA lines). These artifacts were not genuine false positives but rather inconsistencies arising from assembly-based methods, making this criterion unreliable as a validation metric.

*Why the Second and Third Criteria Were Insufficient*

The correlation between SV and SNP diversity (**Supplementary Table S6**) was indeed high, but this alone was not sufficient to rule out a high false positive rate. Even if the pipeline doubled the number of false positives relative to the true signal, the correlation could still appear strong if false positives were distributed evenly across samples. Similarly, while we observed a decline in the rate of novel SV discovery as sample size increased, this trend is also influenced by false positive rates—pipelines with higher false positive rates will still exhibit a declining discovery rate, rendering this criterion insufficient for validation.

**Section 2**

*Lower Signal-to-Noise Ratio in Spontaneous SV Calls*

Spontaneous SV calls required correction far more frequently upon IGV visualization compared to segregating SV calls. This unexpected observation complicates efforts to develop an automated workflow based on segregating SVs as ‘ground truth’ that can be generalized to call SVs in MA lines. The reason for this discrepancy is likely due to differences in SV size distributions.

Most segregating SVs are relatively small: approximately 82% of all deletions among segregating SVs fall within the 30–300 bp range, with nearly 90% measuring under 1 kb. Similarly, about 74% of segregating insertions are smaller than 300 bp, and ~86% are below 1 kb. In contrast, the size distribution of spontaneous SVs exhibits a multimodal pattern with a heavy tail toward larger variants (**Supplementary Figures S1–S3**), with roughly 45% of spontaneous insertion calls exceeding 1 kb.

This size discrepancy suggests that a pipeline trained on segregating variants as ground truth might be well-suited for detecting smaller SVs but may struggle with larger SVs, which are more prevalent in MA lines. A bias toward smaller variants is expected among wild isolates, as larger SVs are more likely to be deleterious.

Even after excluding the outlier MA517, which had 35 false positive calls for 5 true positives, we found that the signal-to-noise ratio for spontaneous SVs in the remaining MA lines was 2:1 at best. This is considerably lower than the signal-to-noise ratio observed for segregating SVs: 19:1 for small deletions under 300 bp (n = 40), 5.6:1 for larger deletions around 1 kb (n = 20), 19:1 for small insertions under 300 bp (n = 40), and 2.3:1 for larger insertions around 1 kb (n = 20).

False positive rates were estimated through IGV visualization of a random sample of 40 deletions and insertions (<300 bp) and 20 larger deletions and insertions (~1 kb). The loci selected for this analysis, along with manual annotations, are provided in **Supplementary Tables S7 and S8** for deletions and insertions, respectively.

**Section 3**

*A Common Limitation of Assemblytics, SMRT-SV2, and Sniffles*

A notable challenge in SV calling arises when a repeating unit is duplicated or deleted. In such cases, alternative alignments of reads or pseudo-reads can create the appearance of heterozygosity within a sample or an apparent mutation across samples. This is a common type of false positive we encountered with tools such as Assemblytics, SMRT-SV2 (default; without the modifications we introduced), and Sniffles. In contrast, PBSV handled this issue far more effectively, likely due to its internal assembly step prior to variant calling. While the inner workings of PBSV remain unclear beyond what is described in the software manual—since it is distributed as a binary by PacBio without an accompanying publication—our results demonstrate that in this particular use case, PBSV vastly outperforms other tools.

One of the other issues with de novo assembly and genome-to-genome alignment (e.g., Assemblytics) is that while it is the most reference-free approach (minimizing reference bias) for SV calling, it is highly dependent on the quality of the assemblies. Poor assemblies across certain loci can create the illusion of SVs wherever one assembly disagrees with another, introducing a major source of false positives. This limitation ultimately guided us toward the SMRT-SV2 architecture (away from Assemblytics), which relies on alignment first, followed by local assembly across tiled regions, providing a more balanced approach that mitigates both reference bias and assembly artifacts.

Ultimately, we resorted to manual inspection of each variant call in the MA lines to evaluate the veracity of the call. Together with a simulation-based estimate of the false negative rate, this workflow is laborious but sufficient to call spontaneous SVs in MA lines.

**Supplementary Methods**

*Estimation of Pairwise Genetic Distance*
To estimate genetic distance based on base substitutions and small indels, we developed custom AWK scripts to process variant call format (VCF) files. A pairwise genetic distance of 1 unit was assigned when both lines were homozygous for different alleles. When one line was heterozygous and the other was homozygous with a shared allele, a genetic distance of 0.5 units was assigned.

*De novo Genome Assembly*

*De novo* genome assembly using Canu (Koren et al. 2017) is computationally intensive, requiring substantial resources. To optimize the assembly, we conducted 88 iterations on the N2 genome, systematically varying multiple parameters, including coverage, sketch size, k-mer size during correction, overlap-based trimming, and assembly settings. Among these, increasing the sketch size yielded the most significant improvements, while varying coverage had no meaningful impact on assembly quality.

We present a summary of all de novo assemblies in **Supplementary Tables S9–S11**, which document the progression of our optimization process. **Table S9** illustrates the initial optimizations, confirming that increasing the sketch size is the most effective strategy. **Table S10** explores the impact of varying k-mer sizes during the correction phase, while **Table S11** evaluates k-mer size adjustments during the overlap stage of Canu.

*Assemblytics pipeline*

Assemblytics is a web-based structural variant detection tool ([*http://assemblytics.com/*](http://assemblytics.com/)) with a Linux command-line interface. The web version allows limited parameter customization, so we used the command-line version for greater flexibility. We retained the default "Unique Sequence Length Required" setting of 10,000 bp as a proxy for mapping quality, where longer unique sequences improve accuracy. The maximum variant size was increased from 10,000 bp to 100,000 bp (the largest permissible value), and the minimum was lowered from 50 bp to 30 bp.

To generate the required delta file, we followed the authors’ recommendations using the following *nucmer* command:

nucmer --maxmatch -t 8 -l 100 -c 500 -p $LINE $REFERENCE $FASTA_FILE

The -l and -c parameters follow the authors' guidelines, while -t enables multi-threading. The resulting delta file, generated using nucmer (Marcais et al. 2018), was compressed with gzip, as recommended, and used as input for Assemblytics.

The output BED file is formatted for bedtools (Quinlan, Hall 2010) and processed using custom BASH scripts to estimate nucleotide diversity (θ) and mutation rate (µ). To identify overlapping variant calls, we used bedtools intersect without imposing a minimum overlap threshold. Since bedtools intersect is not inherently variant-aware, all analyses were conducted separately for each variant type.

*Estimation of variant overlaps*

Variant overlaps were identified using **bedtools intersect** (Quinlan, Hall 2010). Overlaps were assessed using a permissive criterion, recording any instance where at least one base pair overlapped, provided both variants were of the same type.
