## Supplementary Sequence File S1 for "High rate of mutation and efficient removal by selection of structural variants from natural populations of *Caenorhabditis elegans*"

The first sequence, a **3.1 kb insertion**, was extracted from an assembled contig (pseudo-read) of the **PB306 ancestor**. The second sequence, a **1.9 kb insertion**, was obtained from a pseudo-read of the **PB306-derived MA line MA 445**. The final sequence represents a **173-bp repeating motif** that is not characterized by RepeatMasker, meaning the defined start and end points of a standardized repeat may vary.

>3.1kb_insertion

CCGGCAAATTGCCGAAAATGAAAATCTCCGGCAAACCGGCAATTTGCTGATTTGCCGGAAAACTGGCAATTCCCCTCTAAACGTTTCCGGCAAATTTCCAGAGTTGACAATTTCCAGCAAACCGGCAATTTTCTGATTTGCCGGAAAAACGGCAAATTGCCCTCTAAACATTTCCGGCAAATCTGATATCCGGCAAATTGCCGAAAATGAAAATTTCCGGCAAACCGGCAATTTGCTGATTTGCCGGAAAACTGGCAATTCCCCTCTAAACGTTTCCGGCAAATTTCCAGAGTTGACAATTTCCAGCAAACCGGCAATTTTCTGATTTGCCGGAAAAACGGCAAATTGCCCTCTAAACATTTCCGGCAAATCTGATATCCGGCAAATTGCCGAAAATGAAAATTTCCGGCAAACCGGCAATTTGCTGATTTGCCGGAAAACTGGCAATTCCCCTCTAAACGTTTCCGGCAAATTGCCAGAGTTGACAATTTCCAGCAAACCGGCAATTTTCTGATTTGTCGGAAAAACGGCAAATTGCCCTCTAAACATTTCCGGCAAATTGCCAGAGTTGACAATTTCCAGCAAACCGGCAATTTGCTGATTTGCCGGAAAACTGGCAATTCCCCTCTAAACGTTTCCGGCAAATTGCCAGAGTTGACAATTTCCAGCAAACCGGCAATTTTCTGATTTGTCGGAAAAACGGCAAATTGCCCTCTAAACATTTCCGGCAAATTGCCAGAGTTGACAATTTCCAGCAAACCGGCAATTTGCTGATTTGCCGGAAAACTGGCAATTCCCCTCTAAACGTTTCCGGCAAATTGCCAGAGTTGACAATTTCCAGCAAACCGGCAATTTTCTGATTTGTCGGAAAAACGGCAAATTGCCCTCTAAACATTTCCGGCAAATTGCCAGAGTTGACAATTTCCAGCAAACCGGCAATTTGCTGATTTGCCGGAAAACTGGCAATTCCCCTCTAAACGTTTCCGGCAAATTGCCAGAGTTGACAATTTCCAGCAAACCGGCAATTTTCTGATTTGTCGGAAAAACGGCAAATTGCCCTCTAAACATTTCCGGCAAATTGCCAGAGTTGACAATTTCCAGCAAACCGGCAATTTGCTGATTTGCCGGAAAACTGGCAATTCCCCTCTAAACGTTTCCGGCAAATTGCCAGAGTTGACAATTTCCAGCAAACCGGCAATTTTCTGATTTGTCGGAAAAACGGCAAATTGCCCTCTAAACATTTCCGGCAAATTGCCAGAGTTGACAATTTCCAGCAAACCGGCAATTTTCTGATTTGTCGGAAAAACGGCAAATTGCCCTCTAAACATTTCCGGCAAATTGCCAGAGTTGACAATTTCCAGCAAACCGGCAATTTGCTGATTTGTCGGAAAAACGGCAAATTGCCCTCTAAACATTTCCGGCAAATTGCCAGAGTTGACAATTTCCAGCAAACCGGCAATTTGCTGATTTGCCGGAAAAACTGGCAATTCCCCTCTAAACGTTTCCGGCAAATTGCCAGAGTTGACAATTTCCAGCAAACCGGCAATTTTCTGATTTGTCGAAAAACGGCAAATTGCCCTCTAAACATTTCCGGCAAATTGCCAGAGTTGACAATTTCCAGCAAACCGGCAATTTGCTGATTTGCCGGAAAACTGGCAATTCCCCTCTAAACGTTTCCGGCAAATTGCCAGAGTTGACAATTTCCAGCAAACCGGCAATTTTCTGATTTGTCGGAAAAACGGCAAATTGCCCTCTAAACATTTCCGGCAAATTGCCAGAGTTGACAATTTCCAGCAAACCGGCAATTTGCTGATTTGCCGGAAAACTGGCAATTCCCCTCTAAACGTTTCCGGCAAATTGCCAGAGTTGACAATTTCCAGCAAACCGGCAATTTTCTGATTTGTCGGAAAAACGGCAAATTGCCCTCTAAACATTTCCGGCAAATTGCCAGAGTTGACAATTTCCAGCAAACCGGCAATTTGCTGATTTGCCGGAAAACTGGCAATTCCCCTCTAAACGTTTCCGGCAAATTGCCAGAGTTGACAATTTCCAGCAAACCGGCAATTTTCTGATTTGTCGGAAAAAACGGCAAATTGCCCTCTAAACATTTCCGGCAAATTGCCAGAGTTGACAATTTCCAGCAAACCGGCAATTTTCTGATTTGTCGGAAAAACGGCAAATTGCCCTCTAAACATTTCCGGCAAATTGCCAGAGTTGACAATTTCCAGCAAACCGGCAATTTGCTGATTTGTCGGAAAAACGGCAAATTGCCCTCTAAACATTTCCGGCAAATTGCCAGAGTTGACAATTTCCAGCAAACCGGCAATTTGCTGATTTGCCGGAAAACTGGCAATTCCCCTCTAAACGTTTCCGGCAAATTGCCAGAGTTGACAATTTCCAGCAAACCGGCAATTTTCTGATTTGTCGGAAAAACGGCAAATTGCCCTCTAAACATTTCCGGCAAATTGCCAGAGTTGACAATTTCCAGCAAACCGGCAATTTGCTGATTTGCCGGAAAACTGGCAATTCCCCTCTAAACGTTTCCGGCAAATTGCCAGAGTTGACAATTTCCAGCAAACCGGCAATTTTCTGATTTGTCGGAAAAACGGCAAATTGCCCTCTAAACATTTCCGGCAAATTGCCAGATGTTGACAATTTCCAGCAACCGGCAATTTTCTGATTTGTCGGAAAAACGGCAAATTGCCCTCTAAACATTTCCGGCAAATTGCCAGAGTTGACAATTTCCAGCAAACCGGCAATTTGCTGATTTGTCGGAAAAACGGCAAATTGCCCTCTAAACATTTCCGGCAAATTGCCAGAGTTGACAATTTCCAGCAAACCGGCAATTTGCTGATTTGCCGGAAAACTGGCAATTCCCCTCTAAACGTTTCCGGCAAATTGCCAGAGTTGACAATTTCCAGCAAACCGGCAATTTTCTGATTTGTCGGAAAAACGGCAAATTGCCCTCTAAACATTTCCGGCAAATTGCCAGAGTTGACAATTTCCAGCAAACCGGCAATTTGCTGATTTGCGGAAAACTGGCAATTCCCCTCTAAACGTTTCCGGCAAATTGCCAGAGTTGACAATTTCCAGCAAACCGGCAATTTTCTGATTTGTCGGAAAAACGGCAAATTGCCCTCTAAACATTTCCGGCAAATCTGATAT

>1.9kb_insertion

CCGGCAAATTGCCGAAAATGAAAATCTCCGGCAAACCGGCAATTTGCTGATTTGCCGGAAAACTGGCAATTCCCCTCTAAACGTTTCCGGCAAATTTCCAGAGTTGACAATTTCCAGCAAACCGGCAATTTTCTGATTTGCCGGAAAAACGGCAAATTGCCCTCTAAACATTTCCGGCAAATCTGATATCCGGCAAATTGCCGAAAATGAAAATTTCCGGCAAACCGGCAATTTGCTGATTTGCCGGAAAACTGGCAATTCCCCTCTAAACGTTTCCGGCAAATTTCCAGAGTTGACAATTTCCAGCAAACCGGCAATTTTCTGATTTGCCGGAAAAACGGCAAATTGCCCTCTAAACATTTCCGGCAAATCTGATATCCGGCAAATTGCCGAAAATGAAAATTTCCGGCAAACCGGCAATTTGCTGATTTGCCGGAAAACTGGCAATTCCCCTCTAAACGTTTCCGGCAAATTGCCAGAGTTGACAATTTCCAGCAAACCGGCAATTTTCTGATTTGTCGGAAAAACGGCAAATTGCCCTCTAAACATTTCCGGCAAATTGCCAGAGTTGACAATTTCCAGCAAACCGGCAATTTGCTGATTTGCCGGAAAACTGGCAATTCCCCTCTAAACGTTTCCGGCAAATTGCCAGAGTTGACAATTTCCAGCAAACCGGCAATTTTCTGATTTGTCGGAAAAAACGGCAAATTGCCCTCTAAACATTTCCGGCAAATTGCCAGAGTTGACAATTTCCAGCAAACCGGCAATTTGCTGATTTGCCGGAAAACTGGCAATTCCCCTCTAAACGTTTCCGGCAAATTGCCAGAGTTGACAATTTCCAGCAAACCGGCAATTTTCTGATTTGTCGGAAAAACGGCAAATTGCCCTCTAAACATTTCCGGCAAATTGCCAGAGTTGACAATTTCCAGCAAACCGGCAATTTGCTGATTTGCCGGAAAACTGGCAATTCCCCTCTAAACGTTTCCGGCAAATTGCCAGAGTTGACAATTTCCAGCAAACCGGCAATTTTCTGATTTGTCGGAAAAACGGCAAATTGCCCTCTAAACATTTCCGGCAAATTGCCAGAGTTGACAATTTCCAGCAAACCGGCAATTTGCTGATTTGCCGGAAAACTGGCAATTCCCCTCTAAACGTTTCCGGCAAATTGCCAGAGTTGACAATTTCCAGCAAACCGGCAATTTTCTGATTTGTCGGAAAAACGGCAAATTGCCCTCTAAACATTTCCGGCAAATTGCCAGAGTTGACAATTTCCAGCAAACCGGCAATTTTCTGATTTGTCGGAAAAACGGCAAATTGCCCTCTAAACATTTCCGGCAAATTGCCAGAGTTGACAATTTCCAGCAAACCGGCAATTTGCTGATTTGTCGGAAAAACGGCAAATTGCCCTCTAAACATTTCCGGCAAATTGCCAGAGTTGACAATTTCCAGCAAACCGGCAATTTGCTGATTTGCCGGAAAACTGGCAATTCCCCTCTAAACGTTTCCGGCAAATTGCCAGAGTTGACAATTTCCAGCAAACCGGCAATTTTCTGATTTGTCGGAAAAACGGCAAATTGCCCTCTAAACATTTCCGGCAAATTGCCAGAGTTGACAATTTCCAGCAAACCGGCAATTTGCTGATTTGCCGGAAAACTGGCAATTCCCCTCTAAACGTTTCCGGCAAATTGCCAGAGTTGACAATTTCCAGCAAACCGGCAATTTTCTGATTTGTCGGAAAAACGGCAAATTGCCCTCTAAACATTTCCGGCAAATTGCCAGAGTTGACAATTTCCAGCAAACCGGCAATTTGCTGATTTGCCGGAAAACTGGCAATTCCCCTCTAAACGTTTCCGGCAAATTGCCAGAGTTGACAATTTCCAGCAAACCGGCAATTTTCTGATTTGTCGGAAAAACGGCAAATTGCCCTCTAAACATTTCCGGCAAATCTGATAT

>repeating_sequence

TGCCAGAGTTGACAATTTCCAGCAAACCGGCAATTTGCTGATTTGCCGGAAAACTGGCAATTCCCCTCTAAACGTTTCCGGCAAATTGCCAGAGTTGACAATTTCCAGCAAACCGGCAATTTTCTGATTTGTCGGAAAAACGGCAAATTGCCCTCTAAACATTTCCGGCAAAT
